# Effects of chitin and chitosan on root growth, biochemical defense response and exudate proteome of *Cannabis sativa*

**DOI:** 10.1101/2022.10.27.514128

**Authors:** Pipob Suwanchaikasem, Shuai Nie, Alexander Idnurm, Jamie Selby-Pham, Robert Walker, Berin A. Boughton

## Abstract

Fungal pathogens pose a major threat to *Cannabis sativa* production, requiring safe and effective management procedures to control disease. Chitin and chitosan are natural molecules that elicit plant defense responses. Investigation of their effects on *C. sativa* will advance understanding of plant responses towards elicitors and provide a potential pathway to enhance plant resistance against diseases. Plants were grown in the *in vitro* Root-TRAPR system and treated with colloidal chitin and chitosan. Plant morphology was monitored, then plant tissues and exudates were collected for enzymatic activity assays, phytohormone quantification, qPCR analysis and proteomics profiling. Chitosan treatments showed increased total chitinase activity and expression of pathogenesis-related (PR) genes by 3-5 times in the root tissues. In the exudates, total peroxidase and chitinase activities and levels of defense proteins such as PR protein 1 and endochitinase 2 were increased. Shoot development was unaffected, but root development was inhibited after chitosan exposure. No significant effects on plant defense were observed upon chitin treatment. These results indicate that colloidal chitosan significantly promoted production and secretion of plant defense proteins in *C. sativa* root system and could be used as a potential elicitor, particularly in hydroponic scenarios to manage crop diseases.

**Highlight:** Chitosan induces defense protein productions and secretions in the root tissues and exudates of *C. sativa*, offering a potential pathway to enhance plant resistance against fungal attack.

## Introduction

*Cannabis sativa* L. has been widely grown for millennia, serving humankind with a range of benefits (Chandra *et al*., 2017b). There are two major *C. sativa* varieties, medicinal cannabis and industrial hemp, which are differentiated by the amount of tetrahydrocannabinol (THC) content in plant dry weight. Medicinal cannabis (high-THC cannabis) is used for treating inflammation, seizure, nausea, vomiting and spasticity (Slawek *et al*., 2022). Industrial hemp (low-THC cannabis) has a robust fiber, used for making cloth, textiles, rope, yarn, paper, and building blocks (Zimniewska, 2022). Additionally, hemp seed is consumed as a food supplement due to high amount of proteins and beneficial polyunsaturated fatty acids (Callaway, 2004). Due to increasing market demand, cannabis agriculture has grown rapidly in the last decade, with increasing reports of negative impacts to plant cultivation, identified by local farmers and processors (Jerushalmi *et al*., 2020; Bodwitch *et al*., 2021). Pest and fungal attacks are one of the major problems for *C. sativa* production (Wang, 2021). Aphids, flea beetles, hemp borers, spider mites and bollworms are the common pests (McPartland *et al*., 2000) and fungal diseases such as grey mold, root rot, charcoal rot, stem canker and powdery mildew have been frequently recorded (Punja *et al*., 2019). Standard practices to control pests and pathogens have not been well established (Punja, 2021). Synthetic pesticides and fungicides like chlorpyrifos-methyl and fluopyram or chemical agents like hydrogen peroxide (H_2_O_2_) and potassium bicarbonate have been applied in the field, posing a major concern for environmental and consumer safety (Craven *et al*., 2019; Sandler *et al*., 2019).

To avoid using chemicals, a number of natural products have been studied to induce plant defense, prompting plants to be more resistant against biotic stress (Thakur and Sohal, 2013; Yakhin *et al*., 2016). Chitin, chitosan, and their derivatives are one of the compound types that can elicit plant defense responses (Li *et al*., 2020). Chitin is an abundant natural polysaccharide, formed by a β-1,4-linkage of *N*-acetyl-D-glucosamine (*N*-GlcNAc) subunits. It is a main structural component of crustacean shells, insect exoskeletons and fungal cell walls (Younes and Rinaudo, 2015). Chitosan is a deacetylated form of chitin. It is less abundant in nature but can be processed from chitin using chemical reactions (Elieh-Ali-Komi and Hamblin, 2016). Several studies have demonstrated the beneficial effects of chitin and chitosan to promote plant defense. For example, treating tomato fruits with chitin suspensions enhanced the gene expressions and protein productions of superoxide dismutase, peroxidase, catalase and chitinase enzymes, reducing grey mold disease caused by *Botrytis cinerea* (Sun *et al*., 2018). Priming tomato leaves with chitosan solution stimulated callose deposition on plant cell wall and accumulation of defense hormones, leading to a reduction of leaf lesions caused by the same fungal pathogen (De Vega *et al*., 2021). However, most of those discoveries have been made in foliar parts of the plant due to ease of treatment and subsequent sampling. How these elicitors impact root systems is less well understood due to the physical challenges of studying roots and difficulty in investigating secretion of plant products into the surrounding soil (Lopez-Moya *et al*., 2017). We recently developed an *in vitro* plant growth device, Root-TRAPR system, for imaging and sampling root material, making it possible to explore the impact of different environmental inputs on root system (Suwanchaikasem *et al*., 2022).

In this study, *C. sativa* was hydroponically grown in the Root-TRAPR system and treated with colloidal chitin and chitosan. We hypothesized that these elicitors would, as for aboveground material, trigger the overall plant defense responses. Their effects on plant root system have been revealed in other crops, for example, mixing chitin into soil altered microbial and fungal community surrounding the rhizosphere of lettuce (Debode *et al*., 2016) and supplying chitosan into plant growth medium induced release of phytohormones, lipid signaling and phenolic compounds into tomato root exudate (Suarez-Fernandez *et al*., 2020). To advance understanding in this area, a proteomics approach was included in our study to characterize secreted proteins in the exudate. Genes decoded to defense proteins were quantified in root tissues using quantitative real-time PCR (qPCR). In addition, root growth parameters, phytohormone levels and defense enzyme activities were also measured to examine plant responses inclusively.

## Materials and methods

### Chemicals

Chitin powder from shrimp shells (practical grade, product code: C7170) and chitosan (medium molecular weight, product code: 448877) were purchased from Sigma Aldrich, US. Acetonitrile (ACN), methanol, formic acid and trifluoroacetic acid (TFA) were liquid chromatography-mass spectrometry (LC-MS) grade solvents (Thermo Fisher Scientific, US). Ethanol was analytical grade (Chem-Supply, Australia). Deionized water was used in the plant growth experiment. Milli-Q water (Merck Millipore, Germany) was used for sample extraction and all instrumental analyses. Hoagland formulation was prepared as previously described (Suwanchaikasem *et al*., 2022).

### Colloidal chitin and chitosan preparation

Five g of chitin or chitosan powder was weighed in a 500-ml Erlenmeyer flask and added with 50 ml of 85% phosphoric acid. Then, another 50 ml of 85% phosphoric acid was slowly added with continuous stirring. The mixture was incubated at 4 °C overnight. Pre-cooled 500 ml of ethanol was added to dilute the colloid and incubated at 4 °C overnight again. The mixture was filtered through double layer of Whatman No 1 filter paper using Buchner funnel and then washed with water (approximately 3 l) until pH is neutral (approximately 5-6). The retentate of colloidal chitin or chitosan was collected in a 50-ml conical tube and frozen at − 80 °C overnight. Lyophilization was carried out using an Alpha 1-4 LD plus freeze-drier (Christ, Germany). Dried colloidal chitin and chitosan were kept at room temperature. Before use, chitin and chitosan were resuspended in a Hoagland solution in final concentrations of 0.1%, 0.2% and 0.5% w/v. The mixture was sonicated for 20 min to allow dispersion before applying to the plant.

### Plant growth and harvest

Industrial hemp seed cv. Ferimon was kindly provided by Southern Hemp Australia. Seeds were surface sterilized using 70% ethanol and 0.04% sodium hypochlorite and germinated in a Petri dish for three days. Seedlings with tap root 4-6 cm in length were transferred to the Root-TRAPR system and grown in a CMP 6010 growth chamber (Conviron, Canada). Growth condition was 16 h light at 25 °C and 8 h dark at 21 °C with constant 60% relative humidity. After maintenance for eight days in standard Hoagland solution, colloidal chitin and chitosan were added into the root growth chamber at 0.1%, 0.2% and 0.5% w/v concentrations. After eight days of the treatment, shoot and root tissues were harvested and ground using mortar and pestle with liquid nitrogen supply and subjected to phytohormones, enzymatic assays and qPCR analyses. Exudate was collected twice on treatment day before applying chitin and chitosan (pre-exudate) and on the last day of experiment (post-exudate) and kept at −80 °C. Six plants were grown for each treatment.

### Root growth measurement

Root morphology was imaged using the WinRHIZO Arabidopsis 2019 software (Regent Instruments, Canada). Root length and root surface area were calculated using the same protocol as previously described (Suwanchaikasem *et al*., 2022). Briefly, root region was manually assigned, and root were automatically detected by the software based on image contrast where the root is brighter than the background. Manual adjustment was carried out when the software misplaced the root. The sum of all root lengths is the total root length and the root length multiplied by the root diameter is the root surface area. Shoot and root fresh weight (FW) was measured using an analytical balance (Ohaus, US) upon sample collections.

### Phytohormone analysis

Phytohormone contents were measured from shoot and root tissues using a targeted LC-MS/MS method with slight modification from the previous protocol (Suwanchaikasem *et al*., 2022). Approximately 100 mg tissue was weighed in a 1.5-ml microcentrifuge tube and extracted with 400 µl of 70% methanol, supplied with 500 ng ml^-1^ of six internal standards ([^2^H_5_]zeatin, [^2^H_2_]indole-3-acetic acid, [^2^H_7_]cinnamic acid, [^2^H_4_]salicylic acid, [^2^H_6_]abscisic acid and dihydro-jasmonic acid). The mixture was vigorously vortexed and centrifuged at 13,000 ×g for 20 min. Supernatant was collected in a LC-MS glass vial and subjected to the instrumental analysis, where the Triple-Quad 6410 LC-MS machine (Agilent Technologies, US) was equipped with Poroshell 120 EC-C18 column (2.7 µm; 2.1 × 100 mm). Column temperature was 45 °C and injection volume was 5 µl. Mobile phase A and B were 0.1% FA in water and ACN, respectively. Flow rate was 300 µl min^-1^ and LC gradient program was as follows: 80% A (0-2 min), 80-50% A (2-3 min), 50-5% A (3-12 min), 5% A (12-16 min), 5-80% A (16-17 min) and 80% A (17-23 min). Gas temperature was 250 °C with the flow of 13 l min^-1^. Nebulizer was set at 55 psi. Capillary voltage was 5,500 and 4,500 V for positive and negative ionization modes, respectively. Multiple reaction monitoring (MRM) transition, collision energy and polarity were set as follows: zeatin (220.1 → 136.1 *m/z*, 14 eV, positive), indole-3-acetic acid (IAA, 176.1 → 130.1 *m/z*, 10 eV, positive), cinnamic acid (CA, 149.1 → 103.1 *m/z*, 20 eV, positive), methyl-IAA (190.1 → 130.0 *m/z*, 16 eV, positive), salicylic acid (SA, 137.0 → 93.0 *m/z*, 16 eV, negative), abscisic acid (ABA, 263.1 → 153.1 *m/z*, 8 eV, negative), jasmonic acid (JA, 209.1 → 59.0 *m/z*, 8 eV, negative), JA-isoleucine (JA-Ile, 322.1 → 129.9 *m/z*, 24 eV, negative), 12-oxo-phytodienoic acid (ODPA, 291.0 → 164.9 *m/z*, 20 eV, negative), [^2^H_5_]zeatin (225.2 → 137.1 *m/z*, 20 eV, positive), [^2^H_2_]IAA (178.1 → 132.0 *m/z*, 12 eV, positive), [^2^H_7_]CA (156.1 → 109.0 *m/z*, 22 eV, positive), [^2^H_4_]SA (141.0 → 97.1 *m/z*, 16 eV, negative), [^2^H_6_]ABA (269.1 → 159.1 *m/z*, 8 eV, negative) and dihydro-JA (211.1 → 59.0 *m/z*, 12 eV, negative). Each sample was injected three times. The average relative peak area was compared against the standard curve, created from 4-6 dilutions of the standard (10-5,000 ng ml^-1^). Calibration range was varied based on hormone level, detected from the samples.

### Peroxidase and chitinase activities

Peroxidase and chitinase activities were measured from tissues and exudates using the method previously described (Suwanchaikasem *et al*., 2022). Briefly, total proteins were extracted from tissue samples using 100 mM potassium phosphate buffer, pH 6.5. For peroxidase assay, protein extracts were treated with 0.025% H_2_O_2_ and 50 mM guaiacol in a 96-wells microplate. The rate of absorbance change at 470 nm over 3 min was evaluated. For chitinase assay, the extracts were treated with 1% w/v colloidal chitin at 37 °C for 2 h and centrifuged at 8,000 ×g for 10 min to stop the reaction. Sodium borate buffer, pH 8.5 was mixed to adjust pH of the mixtures at 95 °C for 5 min. Acidic dimethylaminobenzaldehyde (DMAB) reagent was added to colorize released *N*-GlcNAc at 37 °C for 20 min. Absorbance was measured at 585 nm using an Enspire Multilabel plate reader (Perkin Elmer, US) and evaluated against *N*-GlcNAc standard curve (50-2,000 nM). Peroxidase and chitinase activities were normalized to FW and root surface area (RSA) for tissue and exudate samples, respectively.

### Preparation of exudate proteins

Exudate solution (approximately 15 ml) was centrifuged at 2,500 ×g, 4 °C for 20 min to remove debris. Supernatant was transferred to an Amicon Ultra-15 ml, 10 kDa molecular weight cutoff (MWCO) device (Merck Millipore, Germany) and centrifuged at 4,000 ×g, 4 °C for 40 min to concentrate exudate proteins. Retentate (approximately 200 µl) was collected in a 1.5-ml microcentrifuge tube. Protein content was measured using Bradford assay. Briefly, 20 µl of protein extract was mixed with 180 µl of five-time diluted Bradford reagent (Bio-Rad, US) in a 96-well microplate. The mixture was incubated at room temperature for 10 min. Absorbance was detected at 595 nm using the plate reader. A bovine serum albumin (BSA) standard curve was created from 0 to 100 µg ml^-1^.

Eight µg of proteins were aliquoted from each sample and dried down using a RVC 2-33 vacuum concentrator (Martin Christ, Germany). The proteins were processed through S-Trap micro spin columns (Protifi, US) using manufacturer’s protocol with slight modification. Forty-six µl of 1× lysis buffer (5% sodium dodecyl sulfate in 50 mM triethylammonium bicarbonate (TEAB) buffer, pH 8.5) was added to resuspend the dried pellet and vigorously vortexed. Two µl of 240 mM Tris(2-carboxyethyl)-phosphine (TCEP) was added and incubated at 55 °C for 15 min to reduce proteins. Then, 4 µl of 500 mM iodoacetamide was added and incubated at 37 °C for 30 min in the dark to alkylate proteins. Six µl of 27.5% phosphoric acid was added to denature proteins, followed by adding 330 µl of binding buffer (100 mM TEAB in 90% methanol). S-Trap micro spin column (Protifi, US) was placed into a 2-ml microcentrifuge tube. Sample was applied to the column and centrifuged at 4,000 ×g for 30 s to trap proteins. Sample was loaded twice (200 µl each time). Washing step was carried out three times by adding 150 µl of binding buffer into the column and centrifuged at 4,000 ×g for 1 min. The column was centrifuged at 4,000 ×g for 2 min and transferred into a new 1.5-ml microcentrifuge tube. Fifty µl of 20 µg ml^-1^ trypsin in 50 mM TEAB buffer was carefully added into the column by avoiding trapping air bubble on top of the membrane. The column was loosely capped to limit evaporation loss and incubated at 37 °C for 18 h to digest proteins. The column was rehydrated with 50 µl of 50 mM TEAB buffer for 30 min at room temperature and centrifuged at 4,000 ×g for 1 min. Peptides were then eluted from the column using three elution buffers, 40 µl of 50 mM TEAB buffer, 0.2% FA and 50% ACN, respectively. Centrifugation was performed each time at 4,000 ×g for 1 min. Three hundred µl of water was added to dilute the peptides. To clean up final peptides, the solution was loaded into an Amicon Ultra-0.5 ml, 30 kDa MWCO device (Merck Millipore, Germany) and centrifuged at 13,000 ×g for 20 min. Filtrate was collected and dried down using the vacuum concentrator. Finally, pellet was resuspended with 40 µl of MS loading buffer (2% ACN in water + 0.05% TFA) and centrifuged at 13,000 ×g for 20 min. Fifteen µl peptide solution was transferred to a LC-MS vial and subjected to proteomics LC-MS/MS analysis.

### Proteomics LC-MS/MS data acquisition

The Nano-ESI-LC-MS/MS system, comprising of Ultimate 3000 RSLC (Thermo Fisher Scientific, US), Acclaim Pepmap RSLC analytical column (C18, 100 Å, 75 μm × 50 cm), Acclaim Pepmap nano-trap column (C18, 100 Å, 75 μm × 2 cm) and Q Exactive Plus Orbitrap mass spectrometer (Thermo Fisher Scientific, US) was used for the proteomics LC-MS/MS analysis. Column temperature was 50 °C. Mobile phase A and B was 0.1% FA + 5% dimethyl sulfoxide (DMSO) in water and ACN, respectively. Injection volume was 6 µl. The trap column was loaded with peptide sample at an isocratic flow of 3% ACN containing 0.05% TFA at 6 µl min^-1^ for 6 min, followed by the switch of the trap column as parallel to the analytical column. LC flow rate was 300 nl min^-1^ and gradient was set as follows: 97% A (0-1 min), 97-77% A (1-30 min), 77-60% A (30-40 min), 60-20% A (40-45 min), 20% A (45-50 min), 20-97% A (50-50.1 min) and 97% (50.1-60 min). MS ionization was in positive mode with the settings of 1.9 kV nano electrospray ionization (nESI) voltage, 70% S-lens radio frequency (RF) and capillary temperature of 250 °C. Full scan MS spectra was set at mass range of 375-1400 m/z, maximum ion trapping time of 50 ms, autogain control target value of 3×10^6^ and resolution of 70,000 at m/z 200. Data dependent acquisition (DDA) mode was used to acquire higher-energy collisional dissociation (HCD)-MS/MS spectra of the top 15 most abundant precursor ions from each full scan MS spectrum. The m/z isolation window of 1.2, autogain control target value of 5×10^4^, 30% normalized collision energy, mass range of 200-2,000 m/z, maximum ion trapping time of 50 ms, and resolution of 17,500 at m/z 200 were used to perform HCD-MS/MS of precursor ions (charge states from 2 to 5). Dynamic exclusion of 30 s was enabled.

### Proteomics data analysis

Raw LC-MS/MS data were processed using MaxQuant version 2.0 software (Tyanova *et al*., 2016a). Protein database was gathered from the NCBI web service, including all *C. sativa* proteins annotated from *C. sativa* draft genome sequence (van Bakel *et al*., 2011). When searching against the database, methionine oxidation and protein N-terminal acetylation were set as variable modifications and cysteine carbamidomethylation was a fixed modification. Trypsin was selected as the digestion method, allowing maximum 2 mis-cleavages. Label-free quantification (LFQ) was applied. Reversed hits, potential contaminants and proteins identified only by site were removed from the identification list. Proteins were rigidly filtered using criteria of unique and razor peptides ≥ 2 and peptide sequence coverage ≥ 8%. Proteins with null LFQ intensity in all samples were discarded. In total, 57 high-confidence protein groups were identified from the dataset and protein identification details are presented in Supplementary Table S1.

To understand characteristics of the identified proteins, online DeepGoWeb version 1.0.3 web portal (Kulmanov and Hoehndorf, 2020) was used to predict their cellular location, molecular function and biological process. Prediction threshold was set at zero. Prediction was based on sequence similarity to the annotated proteins in the UniProt database. The score, ranging from zero to one, indicated possibility of the protein to associate with particular cellular location, molecular function and biological process.

### Quantitative real-time PCR

Based on proteomics results, four genes related to pathogenesis-related (PR) protein 1, endochitinase 2, PR protein R major form-like and thaumatin-like protein 1 were selected for validation of gene expression level in the root tissues. Gene and primers details are supplied in Supplementary Table S2. RNA was extracted from root tissues using a RNeasy Plant Mini kit (Qiagen, Germany) according to the manufacturer’s protocol. RNA was eluted from the spin column using 40 µl of nuclease-free water and RNA concentration was measured using a UV5Nano spectrophotometer (Mettler Toledo, US). A 500 ng RNA was converted to complementary DNA (cDNA) using Superscript II reverse transcriptase (Thermo Fisher Scientific, US) and kept at −20 °C. Before undertaking PCR reaction, cDNA dilution was diluted 3 times in water. Primer efficiency was determined on a test sample across 5 concentrations of 10-fold dilutions. PCR primers with efficiency of 90-110% and standard curve linearity (*r*^*2*^) above 0.98 were used in the analysis. A 10 µl reaction was composed of 1× SYBR Green Supermix (Bio-Rad, US), 0.4 µM forward and reverse primers and 1 µl of cDNA. The amplification was performed using CFX384 Touch Real-Time PCR system (Bio-Rad, US) with initial denaturation of 30 s at 95 °C, followed by 40 cycles of 15 s at 95 °C and 40 s at 60 °C. Melt curve was performed with 0.5 °C increment from 65 °C to 95 °C. Lid temperature was set at 95 °C. Quantitative cycle (Cq) and melt temperature data were analyzed using CFX Manager software version 3.1 (Bio-Rad, US). Relative quantification was calculated according to (Livak and Schmittgen, 2001). *C. sativa* cucumber peeling cupredoxin (XP_030477701.1) was used as a reference gene as it was found across post-exudate samples (Supplementary Table S1). Five replicates were performed per treatment.

### Statistical analysis

One-way ANOVA, followed by Tukey’s *post hoc* analysis was used to test significant difference (*P* < 0.05) across all treatments. Paired T-test was used to test the difference between pre- and post-exudates of each treatment (*P* < 0.05). Linear regression was built within the growth curves of root length and surface area. The regression line of each treatment was individually compared against that of control to indicate growth rate variation. Regression coefficient calculated from the interaction between day and treatment with *P* < 0.05 indicated significant difference of the growth rate between control and particular treatment. Statistical analysis and graph plotting were performed using Minitab Statistical software version 20.3 (Minitab, US).

For proteomics data, statistical analysis was performed on raw LFQ intensity using Perseus version 2.0 software (Tyanova *et al*., 2016b). For pre- and post-exudates, one-way ANOVA with permutation-based false discovery rate (FDR) followed by Tukey’s *post hoc* test was used for testing significant difference (*q*-value < 0.05) across all treatments at each timepoint. Paired T-test was used to compare difference between pre- and post-exudate of each treatment. Online MetaboAnalyst version 5.0 software (Pang *et al*., 2021) was employed to construct principal component analysis (PCA) and hierarchical clustering heatmap. To achieve normal distribution, the data were square root (SQRT) transformed and mean centered for the PCA plot, showing the first and second components in x and y axis, respectively. LFQ intensity of post-exudate samples were log transformed and subjected for hierarchical clustering heatmap analysis using Euclidean distance measure and Ward clustering method.

## Results

### Chitosan affects root development

Changes in root development according to chitin and chitosan treatments were clearly visible from the scanned root images (Fig. 1). In control plants, root developed from 36.83 ± 7.34 mm to 99.71 ± 10.96 mm in length and 6.24 ± 1.15 to 20.34 ± 1.90 cm^2^ in surface area within eight days after treatment (Fig. 2A-B). In chitin treatments, root growth was not significantly impacted by the lower chitin concentrations (0.1% and 0.2%) but final root length and surface area of 0.5% chitin treatment were 56.71 ± 10.64% and 56.65 ± 10.87% of the control, respectively (Fig. 2A-B). As shown in Supplementary Fig. S1, root growth rate in 0.5% chitin treatment (4.30 mm day^-1^) was also significantly slower than control (9.10 mm day^-1^). By contrast, chitosan largely affected root development. Final root lengths of 0.1%, 0.2% and 0.5% chitosan treatments were significantly shorter (*P* < 0.05) than control, which were only 41.12 ± 4.55%, 28.49 ± 1.30% and 31.42 ± 6.78% of the control, respectively (Fig. 2A). Likewise, root surface area of 0.1%, 0.2% and 0.5% chitosan treatments were only 43.37 ± 4.36%, 30.60 ± 2.20% and 38.91 ± 6.16% of the control, respectively (Fig. 2B). Root barely expanded upon chitosan exposures with growth rate less than 1.5 mm day^-1^ (Supplementary Fig. S1).

**Fig. 1.**
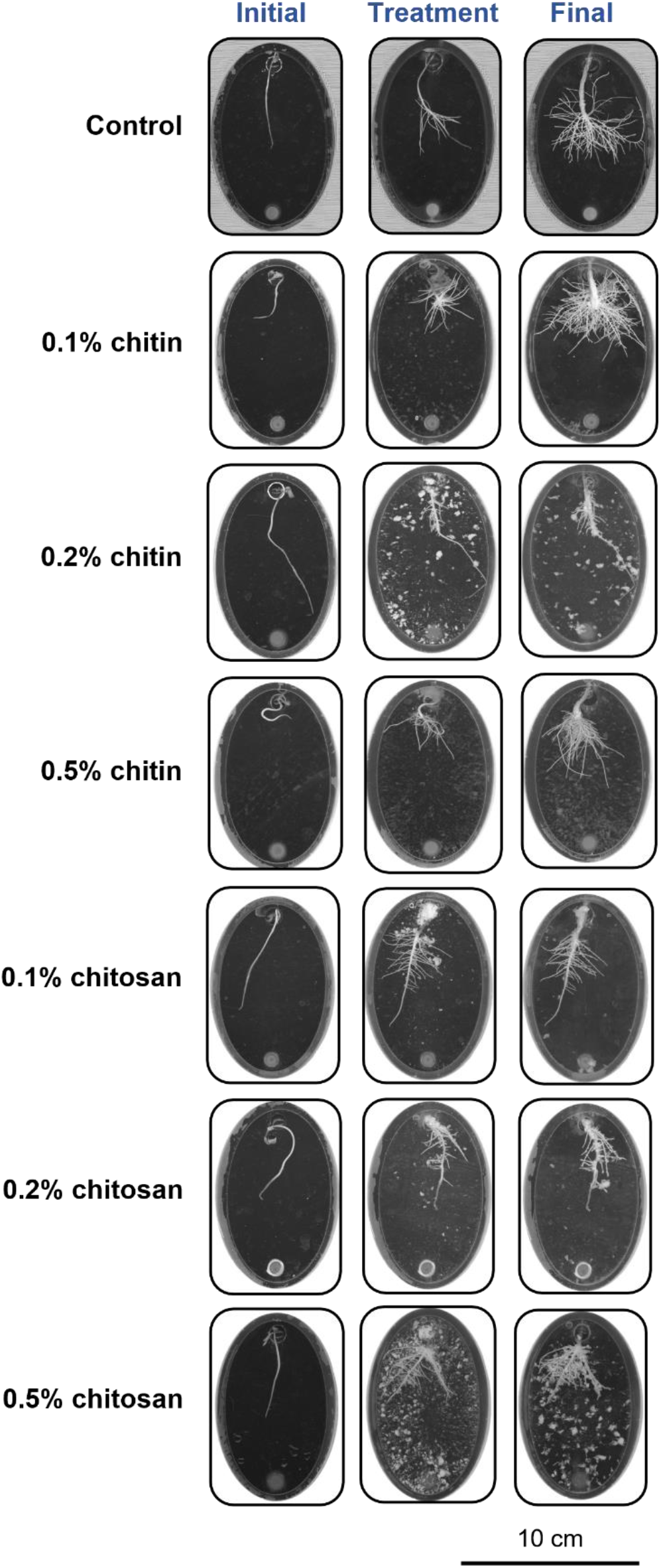
Scanned root images showing root development in the Root-TRAPR system of control, chitin and chitosan treatments. Three scanning times were *initial day* when the seedling was transferred into the Root-TRAPR system, *treatment day (day 0)* when chitin or chitosan was applied into the system and *final day (day 8)* when plant tissues were collected. The images were captured using a calibrated scanner, equipped with the WinRHIZO Arabidopsis 2019 software.

**Fig. 2.**
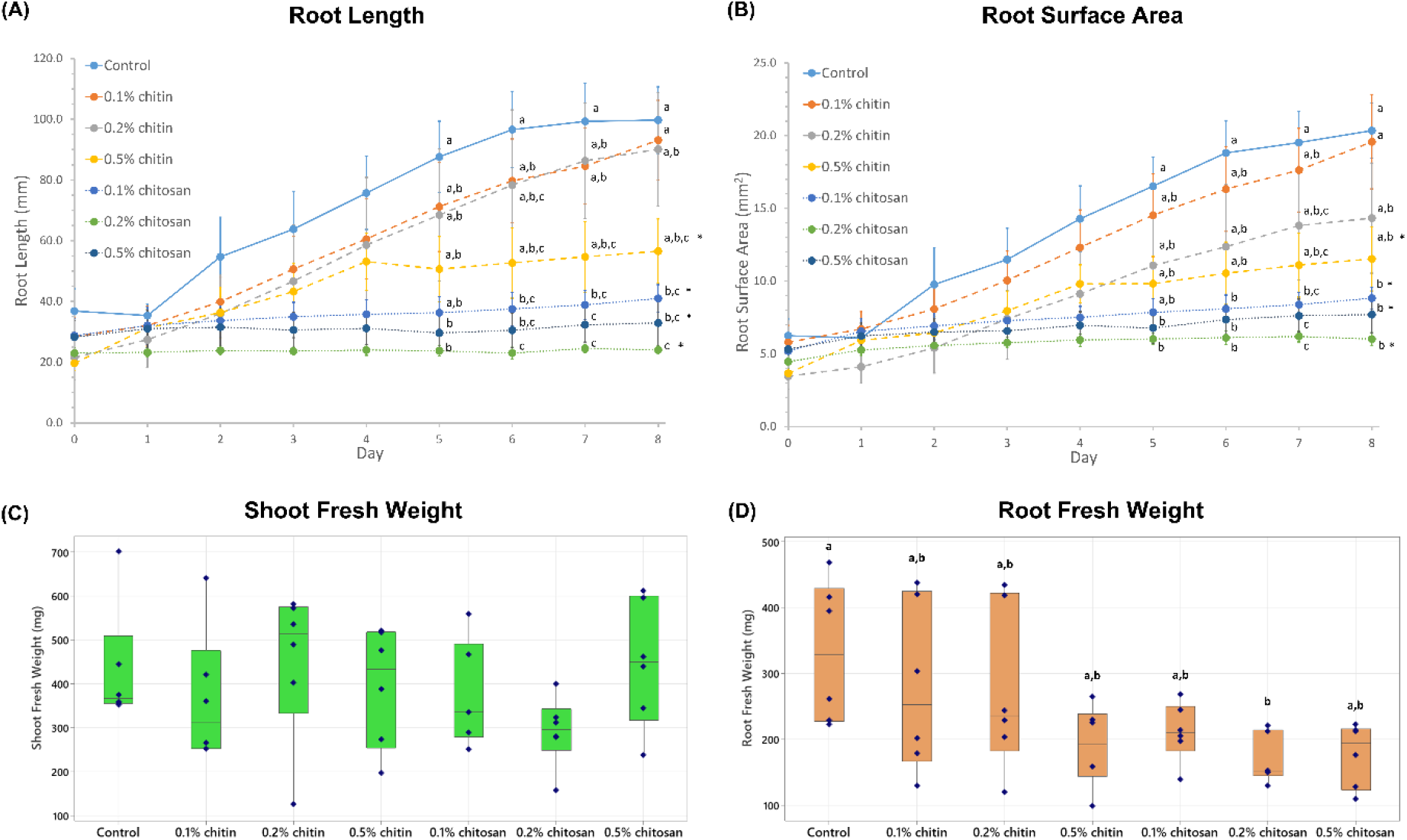
Plant growth measurements showing root length (A), root surface area (B), shoot fresh weight (C) and root fresh weight (D) of control, chitin and chitosan treatments, analyzed from six biological replicates. Chitin and chitosan were applied on day 0 and tissue samples were collected and measured for fresh weight on day 8. Growth curves (A and B) display values of mean ± standard error (SE) bar. Box plots (C and D) display interquartile range box with whiskers and six individual values. Letters (a-c) refer to statistically significant difference (*P* < 0.05) using one-way ANOVA, followed by Tukey’s *post hoc* analysis. Asterisk (*) indicates significant difference (*P* < 0.05) in growth rate, comparing respective treatment with control (Supplementary Fig. S1).

Interestingly, shoot biomass was not impacted by any chitin and chitosan treatments (ANOVA *P* = 0.42), where shoot fresh weights were in the range of 291.52 to 450.97 mg among all treatments (Fig. 2C). Root fresh weight of 0.2% chitosan treatment (168.68 ± 15.42 mg) was significantly lower than control (331.93 ± 43.82 mg). It was approximately 1.5-2 times lower than control in 0.5% chitin (188.97 ± 24.99 mg), 0.1% chitosan (211.07 ± 17.99 mg) and 0.5% chitosan (176.60 ± 19.76 mg) treatments (Fig. 2D).

### Root ABA and CA levels are changed in response to chitosan treatments

The levels of defense hormones including ABA, SA, JA, JA-Ile and OPDA and a plant growth regulator, CA in shoot and root tissues are shown in Fig. 3. The levels of growth hormones including IAA, methyl-IAA and zeatin are presented in Supplementary Fig. S2. In shoot tissues, all hormone levels were relatively comparable among all treatments except methyl-IAA which was significantly lower than control in 0.1% and 0.5% chitosan treatments. In root tissues, ABA and CA levels in 0.5% chitosan treatment (ABA; 36.83 ± 6.41 and CA; 77.05 ± 16.94 ng g^-1^ FW) were significantly higher than those of control (ABA; 14.18 ± 2.48 and CA; 36.29 ± 6.04 ng g^-1^ FW). Moreover, increasing tendencies were observed from SA and OPDA in chitosan treatments but with no significant difference. The levels of SA in 0.2% and 0.5% chitosan treatments (80.97 ± 19.63 and 99.17 ± 29.67 ng g^-1^ FW, respectively) were approximately 2 times higher than control (39.19 ± 6.10 ng g^-1^ FW). Likewise, the OPDA levels of 0.2% and 0.5% chitosan treatments (13.87 ± 2.79 and 14.13 ± 3.67 µg g^-1^ FW, respectively) were nearly 3 times higher than control (5.52 ± 0.83 µg g^-1^ FW).

**Fig. 3.**
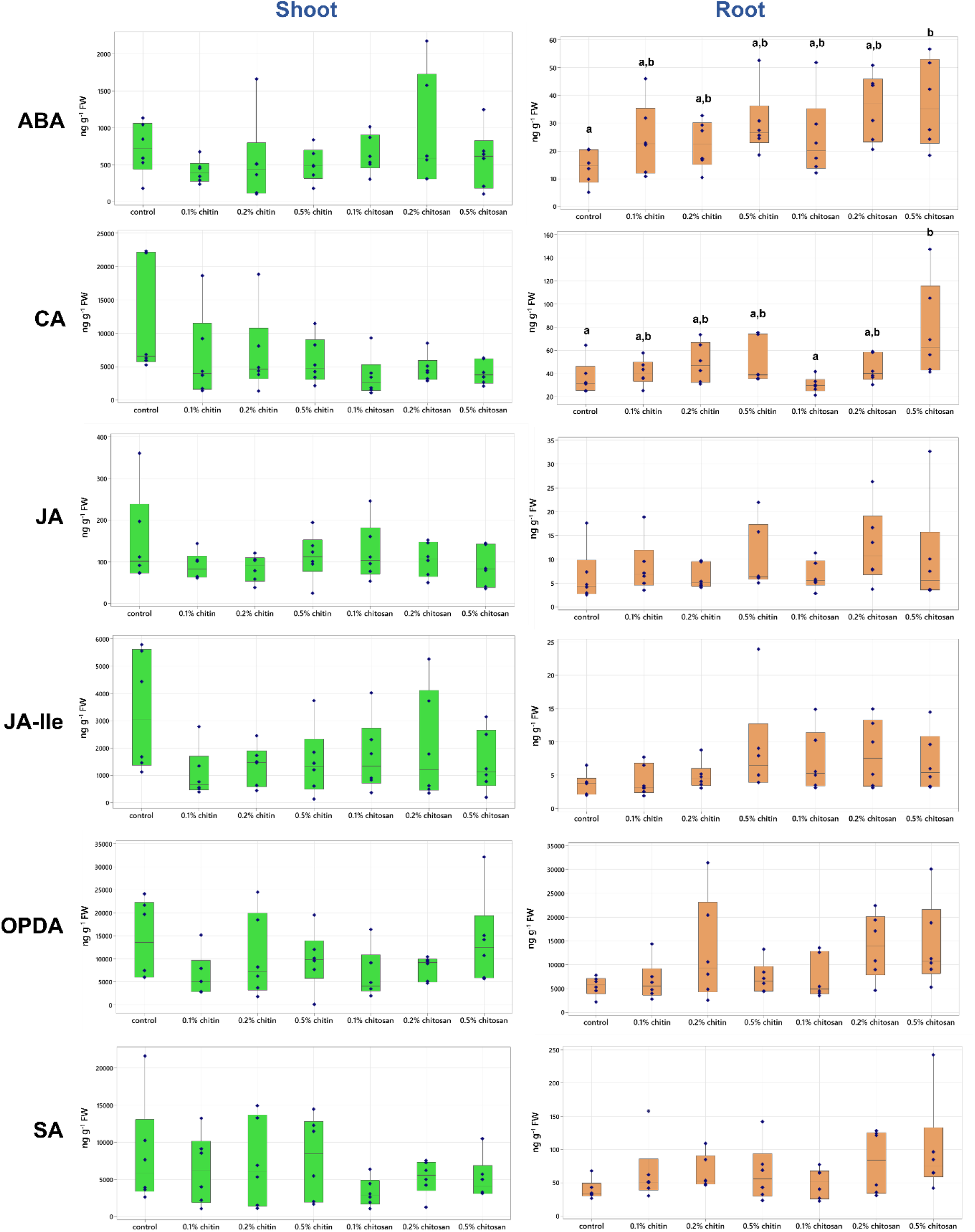
Phytohormone levels of abscisic acid (ABA), cinnamic acid (CA), jasmonic acid (JA), JA-isoleucine (JA-Ile), 12-oxo-phytodienoic acid (OPDA) and salicylic acid (SA) measured from shoot and root tissues of control, chitin and chitosan treatments within six biological replicates. Letters (a-b) refer to statistically significant difference (*P* < 0.05) using one-way ANOVA, followed by Tukey’s *post hoc* analysis.

### Total peroxidase and chitinase activities are increased in root tissues and exudates upon chitosan treatments

Total activities of two plant defense enzymes, peroxidase and chitinase, were measured in shoot, root tissues and exudates. In shoot tissues, peroxidase and chitinase activities were comparable among all treatments (Fig. 4). In root tissues, chitinase activity of 0.2% chitosan treatment (1,880.29 ± 235.16 nmol GlcNAc released g^-1^ FW) were significantly higher than control (678.29 ± 113.42 nmol GlcNAc released g^-1^ FW) but peroxidase activities were comparable across all treatments (837.85 ± 127.48 to 1,124.55 ± 36.85 ΔAbs_470_ min^-1^ g^-1^ FW).

**Fig. 4.**
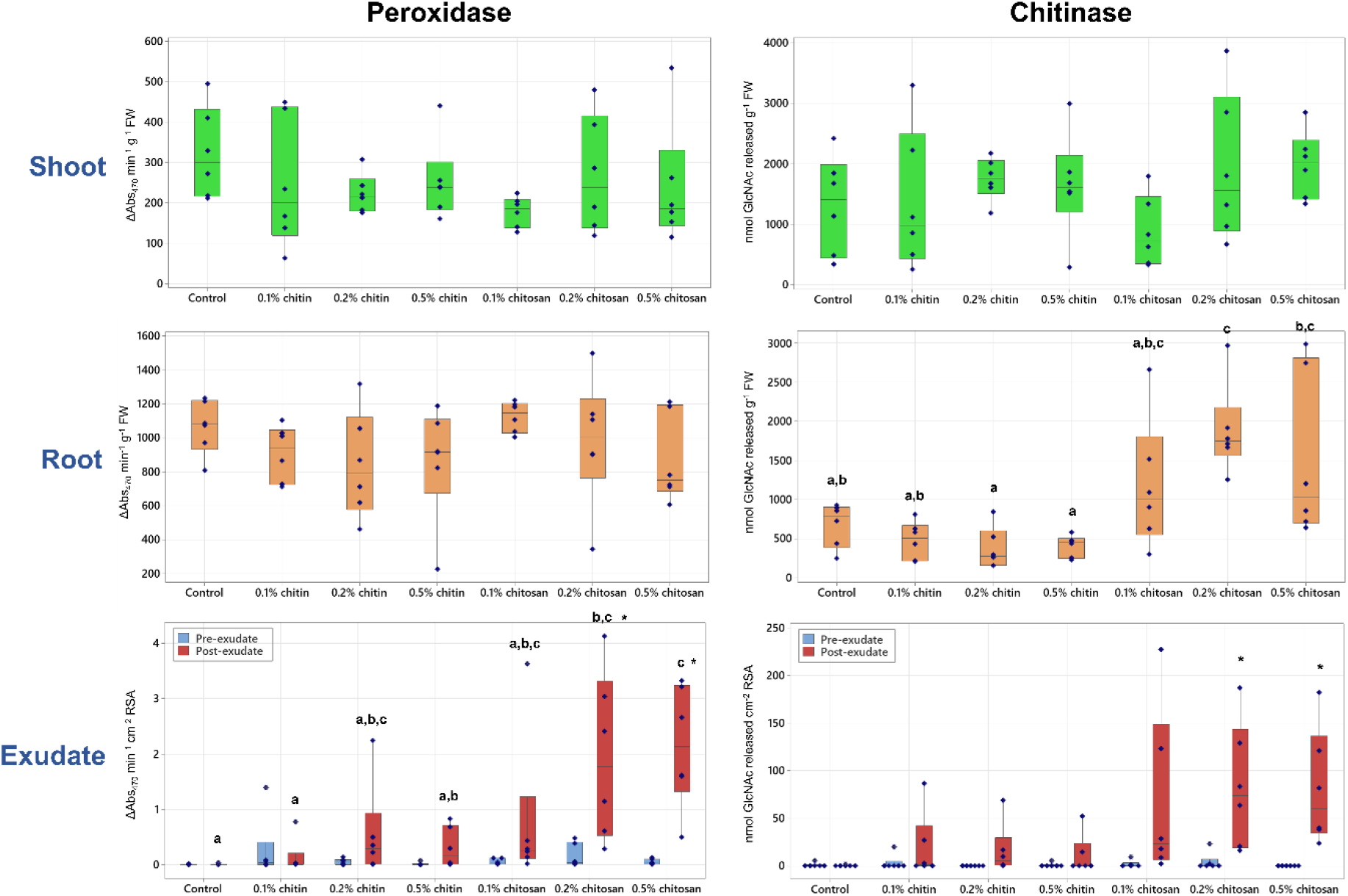
Peroxidase and chitinase activities measured from shoot and root tissues and pre- and post-exudates of control, chitin and chitosan treatments within six biological replicates. Letters (a-c) refer to statistically significant difference (*P* < 0.05) using one-way ANOVA, followed by Tukey’s *post hoc* analysis and asterisk (*) indicates statistically significant difference (*P* < 0.05) between pre- and post-exudate of each treatment using paired T-test.

In exudates, peroxidase and chitinase activities were tested before and after the treatments. Prior to treatment (pre-exudate), peroxidase and chitinase activities were nearly undetectable in all treatments (Fig. 4). After treatment (post-exudate), peroxidase activity of 0.2% and 0.5% chitosan treatments (1.94 ± 0.67 and 2.15 ± 0.49 ΔAbs_470_ min^-1^ cm^-2^ RSA, respectively) was significantly higher than control (0.012 ± 0.006 ΔAbs_470_ min^-1^ cm^-2^ RSA) and their pre-exudates (0.167 ± 0.095 and 0.052 ± 0.023 ΔAbs_470_ min^-1^ cm^-2^ RSA, respectively). Total chitinase activities of all chitosan treatments were substantially increased but not significantly different from control. They were 67.72 ± 40.25, 83.26 ± 29.56 and 81.02 ± 27.41 nmol GlcNAc released cm^-2^ RSA in the 0.1%, 0.2% and 0.5% chitosan treatments, respectively. It was only 0.208 ± 0.208 nmol GlcNAc released cm^-2^ RSA in control. When comparing between before and after treatment, chitinase activities were significantly increased in 0.2% and 0.5% chitosan treatments (Fig. 4). They were 1.79 ± 1.47 and 0.00 ± 0.00 GlcNAc released cm^-2^ RSA in pre-exudates of 0.2% and 0.5% chitosan treatments, respectively and increased to 83.26 ± 29.56 and 81.02 ± 27.41 GlcNAc released cm^-2^ RSA after the treatments. Considering chitin treatments, both peroxidase and chitinase activities tended to increase in post-exudates as compared to control and their pre-exudates. For example, peroxidase activity of post-exudate of 0.2% chitin treatment (0.56 ± 0.38 ΔAbs_470_ min^-1^ cm^-2^ RSA) was approximately 10 times higher than that of pre-exudate (0.056 ± 0.024 ΔAbs_470_ min^-1^ cm^-2^ RSA). Likewise, its chitinase activity was 16.03 ± 11.86 nmol GlcNAc released cm^-2^ RSA, which was totally undetectable in the pre-exudate. However, the effects from chitin treatments were much lesser than chitosan treatments in promoting defense enzyme activities in exudates.

### Chitosan induces root secretion of defense proteins into exudate

Across all samples, 57 protein groups were identified from the root exudates. They were assigned confidently as the identification criteria were restricted to ≥ 2 unique and razor peptides and ≥ 8% sequence coverage. Details of protein identification are found in Supplementary Table S1. PCA was performed to characterize any overall difference of exudate proteomes across all samples. The plot from all samples showed that most pre-exudates (except a small number of outliers: two replicates of 0.1% chitosan, one replicate of 0.1% chitin and one replicate of control) were clustered together, indicating close similarity in their proteome profiles before the treatments (Supplementary Fig. S3A). In post-exudates, all chitin samples were aligned close to the control (Fig. 5A). For chitosan treatment groups, two of the six replicates of 0.1% and 0.2% chitosan treatments were scattered away from the main cluster of control and chitin samples (Fig. 5A-B). All six replicates of 0.5% chitosan were grouped together and clearly segregated from the main cluster, showing clear difference in their proteome profiles after the treatment (Fig. 5B and Supplementary Fig. S3B). The PCA results highlight that exudate proteome profiles were changed according to chitosan treatments, whereby the clearest difference was observed in 0.5% chitosan treatment. To identify the proteins changing upon treatment, hierarchical clustering was applied and a heatmap was generated for post-exudate samples (Fig. 6A). Two major clusters were identified on the horizontal axis, where chitin treatments and control were grouped together, and chitosan treatments were grouped separately. The clustering on the vertical axis, based on relative protein response, displayed three major clusters. The proteins affected by higher concentrations of chitosan (0.2-0.5%) were clustered on the top half of the heatmap. The proteins specifically found in 0.1% chitosan were clustered in the middle, while the proteins relative to control and chitin samples were grouped on a separate branch at the bottom (Fig. 6A). The heatmap results highlight that most of the exudate proteins (approximately 40 proteins of total 57 identified proteins) were increasingly secreted upon 0.1-0.5% chitosan treatments.

**Fig 5.**
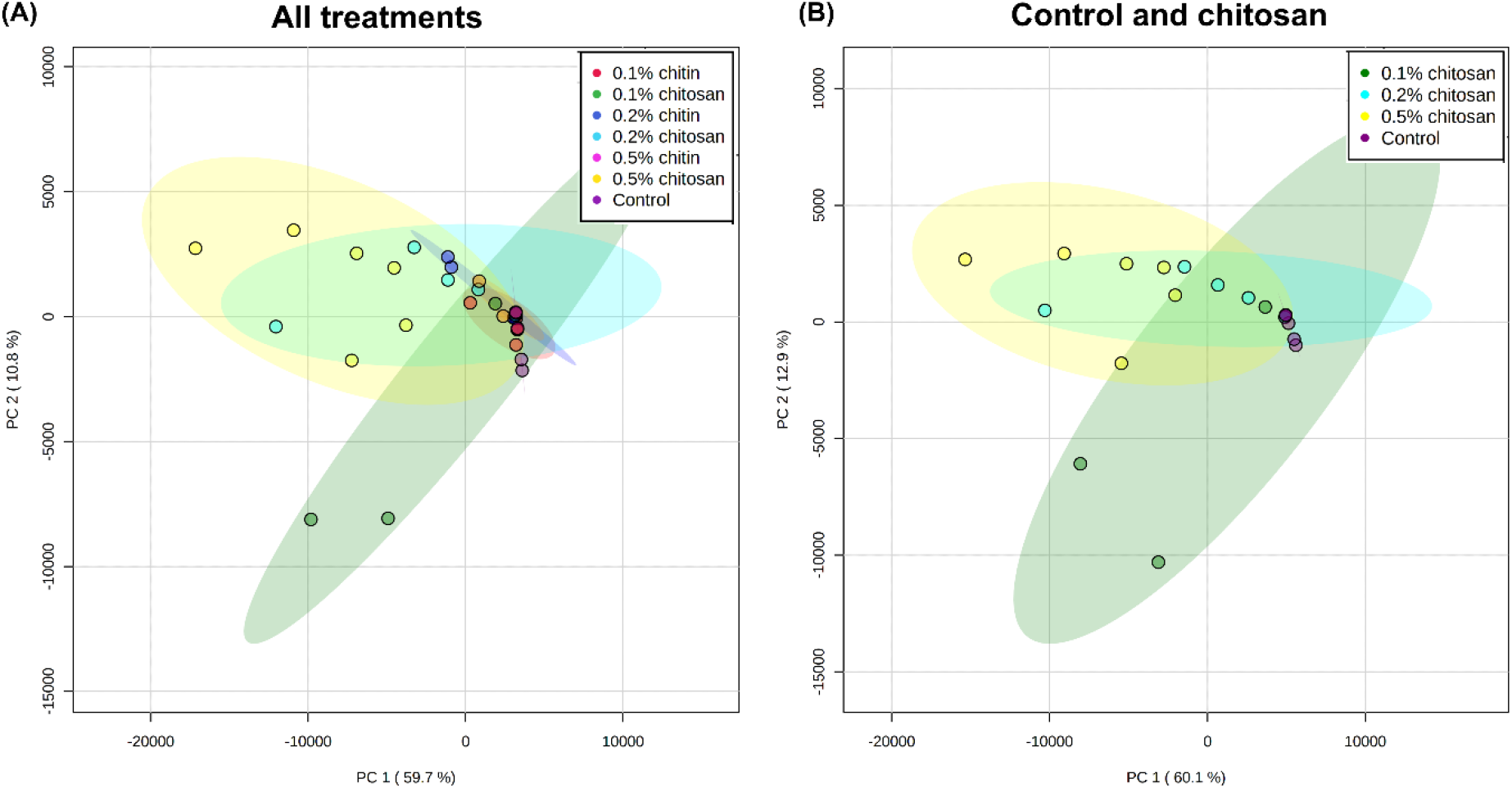
PCA plots of post-exudate proteomes across all control, chitin and chitosan treatments (A) and selectively between control and chitosan treatments (B). Six biological replicates were analyzed per treatment. Control and chitin samples were clustered close together but two replicates of 0.1% chitosan and 0.2% chitosan and six replicates of 0.5% chitosan were clearly separated from the major assembly.

**Fig. 6.**
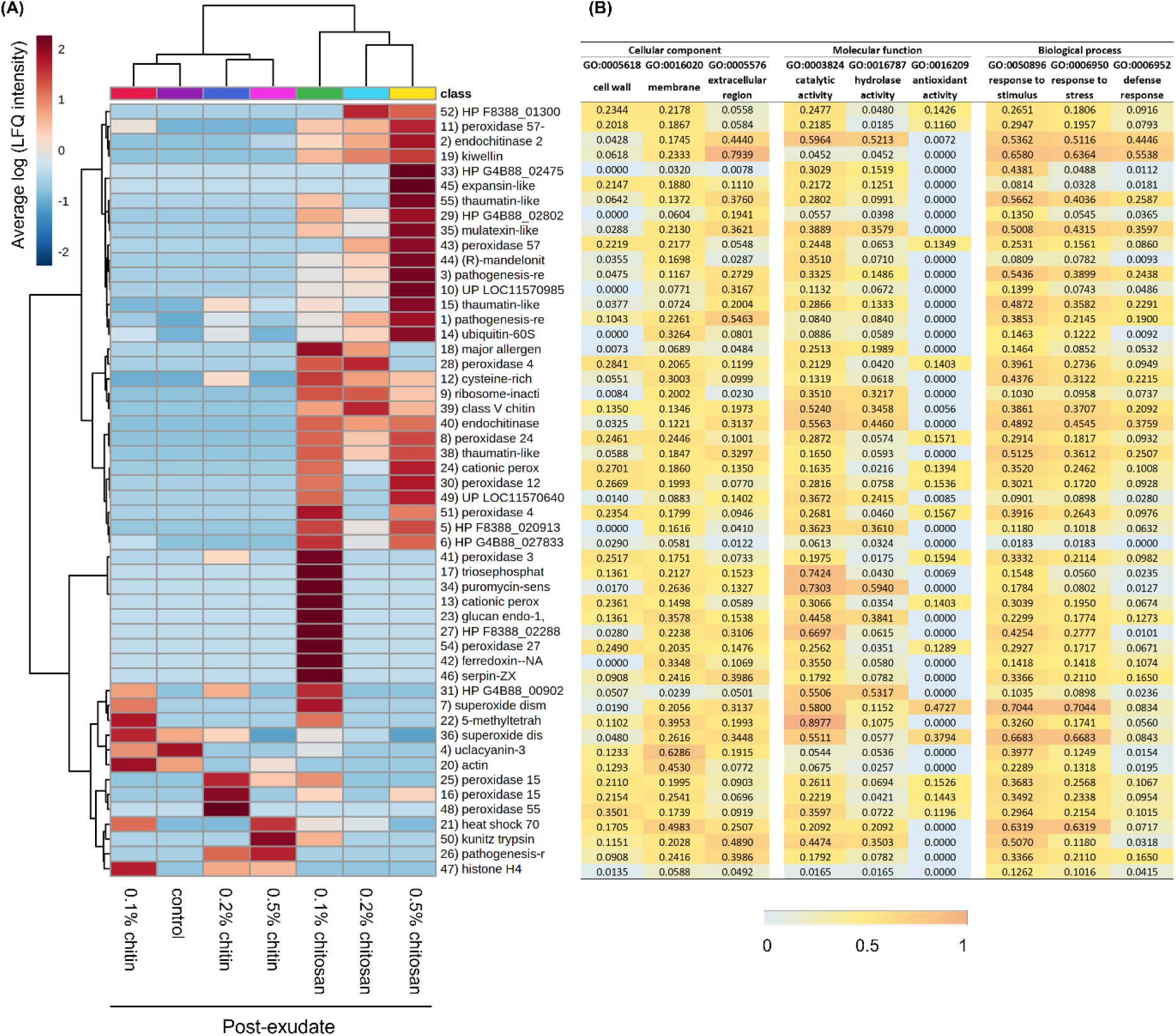
Proteomics analysis displaying clustering heatmap (A) and prediction scores (B) of the exudate proteins. Heatmap shows average of log transformed LFQ intensity within six biological replicates per treatment (A). On horizontal axis, control and chitin samples were clustered together while chitosan treatments were grouped on a separate branch. On vertical axis, proteins highly abundant in 0.2-0.5% chitosan treatments were clustered on the top half of the heatmap. Proteins abundant in 0.1% chitosan were branched in the middle and proteins abundant in control and chitin treatments were clustered at the bottom. Protein number is correlated to protein identification details listed in Supplementary Table S1. Relative prediction scores are presented in three main categories of cellular location, molecular function and biological process (B). From zero to one, higher scores demonstrate an increased possibility of the protein to associate with that particular location, function and process.

Protein function prediction was carried out based on full protein sequence using the DeepGoWeb online webserver. The results are presented in Fig. 6B and Supplementary Table S3 with relative score ranging from zero to one. Higher scores indicate an increased probability of the protein to associate with particular subcellular location, molecular function or biological process. Overall, several extracellular proteins such as PR protein 1, endochitinase 2, mulatexin-like and kiwellin, were identified from the dataset. Their prediction scores were relatively high in the category of extracellular region but low in intracellular categories of cytoplasm, organelle and nucleus (Supplementary Table S3). Peroxidase proteins predicted with high scores in the categories of cell wall, membrane and cell periphery were also detected in the dataset as abundant. This is reasonable because the plant constantly sloughs root end cap cells, leading to a diffusion of cell wall and membrane proteins into the exudate (Badri and Vivanco, 2009; Dubrovskaya *et al*., 2017). Only a few organelle or nucleolar proteins, were detected. This included histone H4, ubiquitin-60S ribosomal protein L40 and heat shock 70 kDa protein-like (Supplementary Table S3). These intracellular proteins were also identified from the root cap secretome of pea (Wen *et al*., 2007), suggesting they may not be uncommon proteins in extracellular region. Intracellular proteins might be derived from sloughed or broken cells and dispersed into root exudate. Overall, the result indicates the effectiveness of our sample preparation protocol and proteomics analysis to isolate and identify root exudate proteins.

Based on protein functions, half of the detected proteins are enzymes such as chitinase, peroxidase, superoxide dismutase and glucosidase (Fig. 6B and Supplementary Table S3). Interestingly, chitinase enzymes including endochitinase 2 and class V chitinase were identified only from chitosan treatments and classified on the top half of the heatmap (Fig. 6A-B). Conversely, superoxide dismutase enzymes were found associated with control and chitin treatments, clustered at the bottom of the heatmap (Fig. 6A-B). Among a variety of peroxidase isoforms, peroxidase 4, peroxidase 57 and peroxidase 24 were found in response to chitosan treatments. Peroxidase 15 and peroxidase 55 were likely to be associated with chitin treatments, but surprisingly none of peroxidase were detected from control (Fig. 6A-B). In terms of biological process (Fig. 6B), most of the enzymes had high prediction scores in the categories of stimulus and stress responses (> 0.3000). For example, two isoforms of endochitinase 2 were predicted with 0.5362 and 0.4892 scores in the category of response to stimulus. The scores were 0.3332 and 0.7044 for peroxidase 3 and superoxide dismutase [Cu-Zn], respectively. However, only chitinase enzymes had relatively high score in the defense response category, where two isoforms of endochitinase 2 were predicted with 0.4446 and 0.3759 scores. The defense response scores of peroxidase 3 and superoxide dismutase [Cu-Zn] were lower than endochitinase 2, which were only 0.0982 and 0.0834, respectively. Since chitinases were increasingly secreted upon chitosan treatments, the results indicate that chitosan has potential to induce root secretion of defense enzymes into exudate. Additionally, non-enzymatic proteins with relatively high prediction score in the category of defense response, for example PR protein R major form-like (0.2438), thaumatin-like protein 1b (0.2587), mulatexin-like (0.3597) and kiwellin (0.5538) were also predominantly detected from chitosan treatments (Fig. 6A-B), This emphasizes the effect of chitosan to promote defense protein and enzyme secretions.

Statistical analysis was then applied to identify proteins of different levels in the exudates. Two analytical aspects were performed: 1) testing across all conditions within pre- or post-exudates using one-way ANOVA and 2) comparing between pre- and post-exudates of each condition using paired T-test. In pre-exudates, there were no significantly different proteins among all studied groups (Supplementary Table S1), suggesting the exudate proteomes of all samples were similar before the treatment. In post-exudates, there were eight significant proteins detected, with seven proteins found to be highly secreted in 0.5% chitosan condition and one protein, uclacyanin-3, found to be significantly higher in control (Fig. 7A and Supplementary Table S1). Comparing between pre- and post-exudates, twelve proteins were significantly increased according to 0.5% chitosan treatment and one protein, endochitinase 2, was significantly higher in 0.2% chitosan condition (Fig. 7B and Supplementary Table S1). Interestingly, most of the significant proteins are well-known PR proteins, such as PR protein 1, PR protein R major form-like, endochitinase 2, thaumatin-like protein 1 and mulatexin-like protein, which generally function once plant experiences pest and pathogen attack (Agrios, 2005; Ferreira *et al*., 2007). However, none were detected as a significant protein in control, 0.1-0.5% chitin and 0.1% chitosan treatments. This data highlights that the root exudate proteome was significantly changed upon chitosan treatments, but this phenomenon was not observed in chitin treatments.

**Fig 7.**
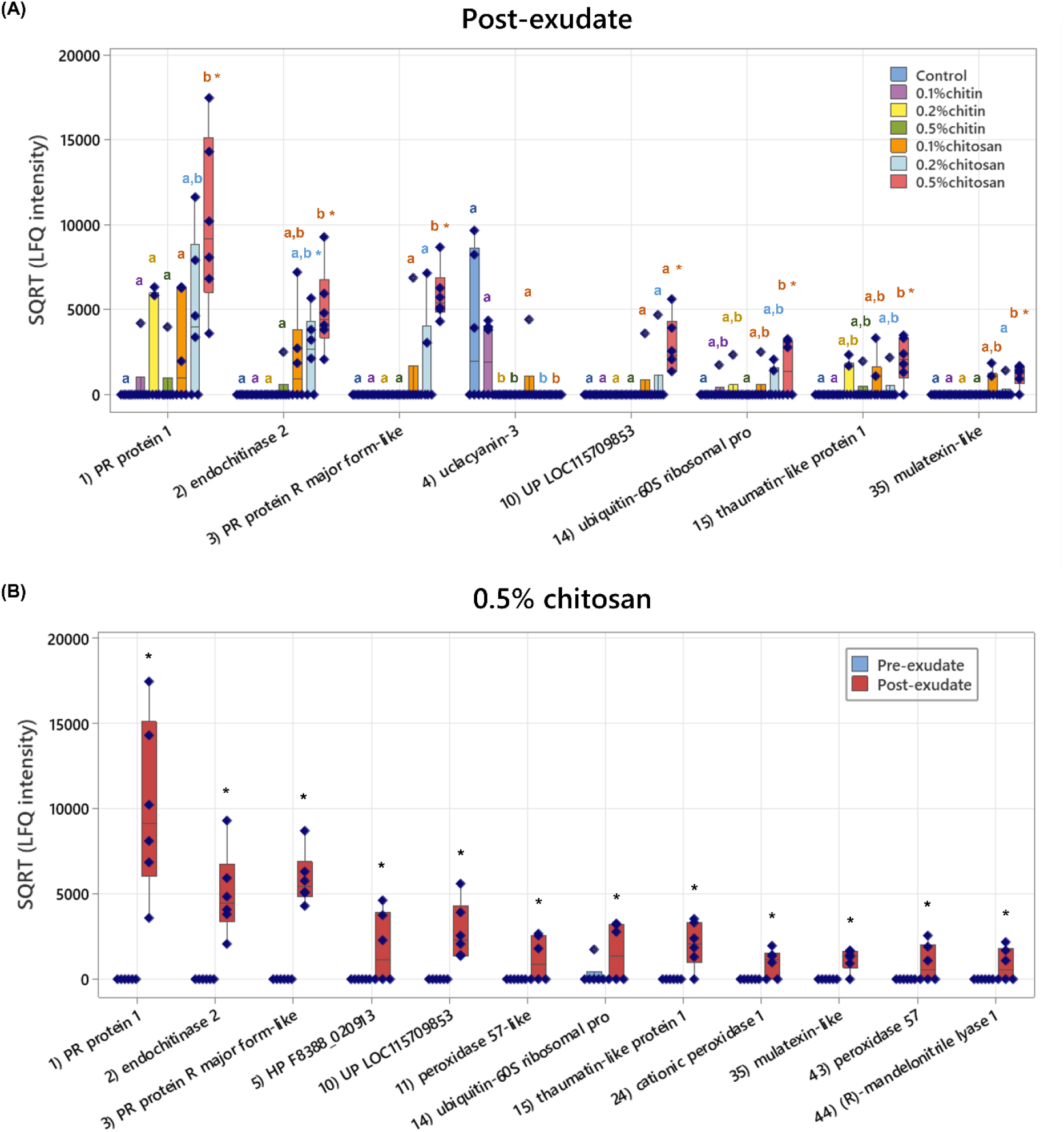
Boxplots of eight significant proteins in the post-exudates across all sample groups (A) and twelve significant proteins between pre- and post-exudates of 0.5% chitosan treatment (B). The plots display interquartile range box with whiskers and individual values within six biological replicates. Letters (a-b) show significant difference (*P* < 0.05) using ANOVA test, followed by Tukey’s *post hoc* analysis and asterisk (*) indicates significant difference (*P* < 0.05) using paired T-test.

### Defense genes are upregulated in root tissues of chitosan-treated plants

Based on exudate proteomics results, four significant defense-related proteins including PR protein 1, endochitinase 2, PR-R major form-like and thaumatin-like protein 1, were selected for qPCR analysis to verify exudate proteome data. Details of the selected genes are described in Supplementary Table S2. Their transcript levels were quantified in the root tissues, collected on the last day of observation. Cucumber peeling cupredoxin was used as a reference gene since it was mostly detected across all the exudate samples (Supplementary Table S1 and S2). It is a copper protein, found in the cell membrane and typically plays a role in electron transport and energy production (Zhang *et al*., 2021).

Relative transcript levels of PR protein 1 were approximately 4.5-times higher than control in the root samples of 0.2% and 0.5% chitosan treatments but only 0.48-1.77 times in other treatments (Fig. 8). Likewise, transcript levels of endochitinase 2 were 2.73 and 2.99 times higher than control in 0.2% and 0.5% chitosan samples, respectively, but only 1.24-1.97 times different in other treatments. The levels of PR protein R major form-like gene were more than two-times higher than control in 0.2% and 0.5% of both chitin and chitosan treatments. The level of thaumatin-like protein 1 was 2.48-times higher than control in 0.2% chitosan treatment. Overall, transcript levels of defense genes tended to increase in root tissues of chitosan-treated samples, but the increases were not significantly different from control. It could be because gene expression processes take place at the early stage once the plant was initially exposed to chitosan and decline over time (Lopez-Moya *et al*., 2017; De Vega *et al*., 2021) and our analysis was performed after the peak period of transcriptome changes had passed. Despite statistically insignificant differences, chitosan-induced gene expressions of PR protein 1, endochitinase 2 and PR protein R major form-like were likely to be dose dependent since positive linear relationships were observed from the plots between qPCR relative fold changes and chitosan concentrations (Supplementary Fig. S4).

**Fig. 8.**
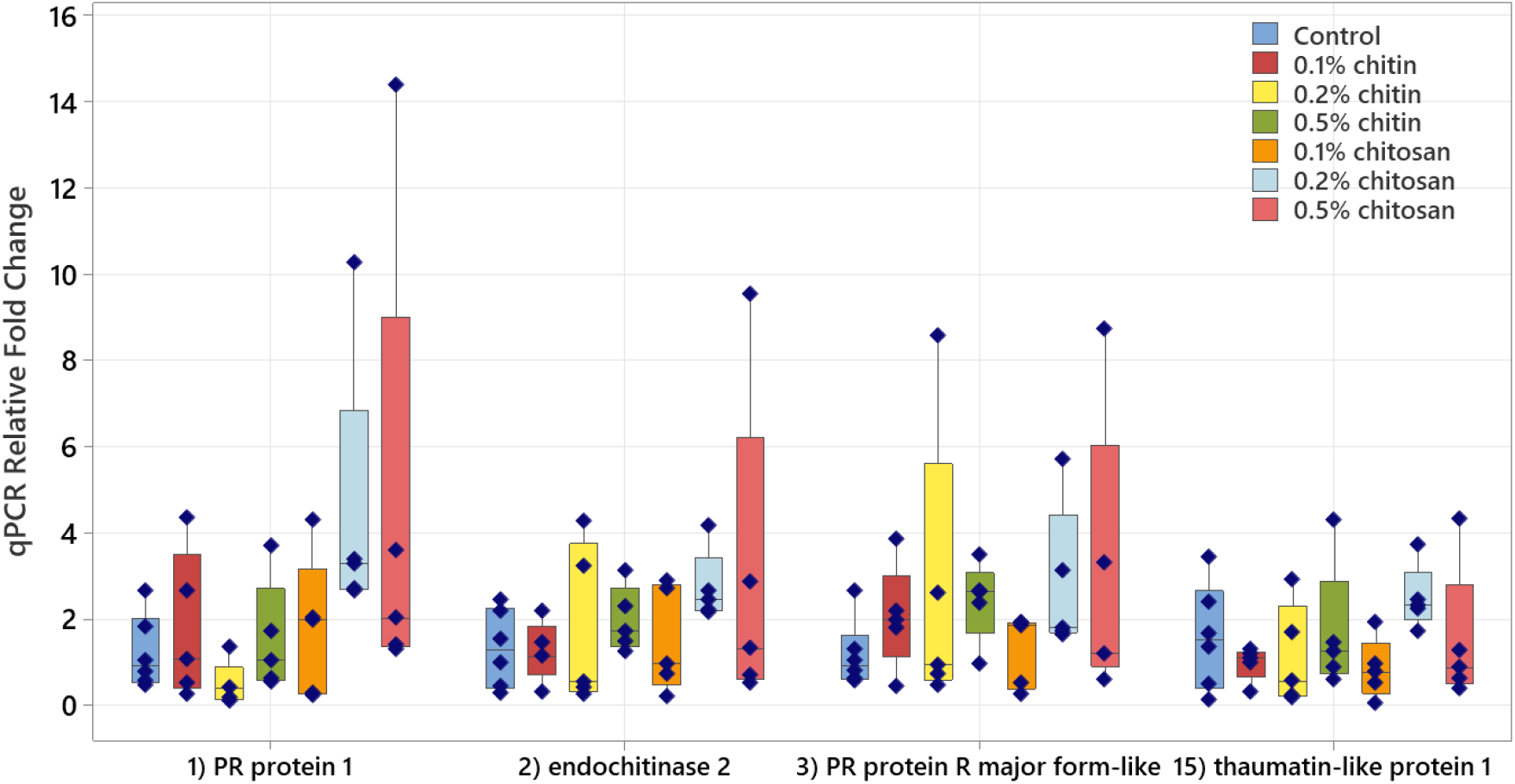
Relative transcript levels of four defense-related genes, encoding PR protein 1, endochitinase 2, PR protein R major form-like and thaumatin-like protein 1, measured from root tissues of chitin and chitosan treatments as normalized to control within five biological replicates.

Furthermore, qPCR data was compared with exudate proteomics and bioassay results. One outlier sample of 0.5% chitosan treatment was identified and removed from the correlation analyses. Positive correlation between endochitinase 2 transcript level and its exudate protein intensity was observed from 0.2% and 0.5% chitosan treatments (Supplementary Fig. S5). It was also detected from the other proteins, PR protein 1, PR-R major form-like and thaumatin-like protein 1, within 0.5% chitosan treatment. Positive correlation was not observed in the lowest chitosan concentration (0.1%) of any proteins (Supplementary Fig. S5). This implies that upon higher doses of chitosan treatment (0.2-0.5%), upregulation of defense gene in the root tissues is likely to contribute to the secretion of specific proteins in the exudates. By contrast, endochitinase 2 transcript level was negatively correlated with total chitinase activities measured from root tissues and exudates (Supplementary Fig. S6), suggesting that total chitinase activities were not influenced by a single chitinase protein but potentially co-contributed by other chitinases in tissue and exudate.

## Discussion

This study aimed to investigate the effects of chitin and chitosan to promote overall defense responses of the *C. sativa* root system. The results demonstrate that chitosan has a much stronger effect than chitin to activate plant root defense responses. Chitosan can induce the production of defense enzymes in root tissue (Fig. 4) and the secretion of defense proteins into root exudate (Fig. 6-7). The gene transcript levels of defense proteins were also upregulated (Fig. 8), and cellular levels of ABA and CA were increased in root tissues (Fig. 3). However, the levels of defense hormones including SA and JA were not changed. In contrast, none of the defense responses induced by chitosan were observed in chitin treatments.

In general, chitin and chitosan may be considered relatively similar in structure. Both are natural β-1,4-linked polysaccharides, with the predominant subunit of *N*-acetyl-D-glucosamine and D-glucosamine for chitin and chitosan, respectively. The difference is the absence of the acetyl group in chitosan subunits. Chitin may be transformed to chitosan by natural chitin deacetylase enzymes (Zhao *et al*., 2010) or chemical deacetylation reactions (Vicente *et al*., 2021). However, the conversion may not be fully complete, and chitosan may be present in a partially acetylated form. The degree of deacetylation determines the number of acetyl groups removed from the chitosan structure. This value along with crystallinity, viscosity and polymer size affect the chemical and physical properties of chitosan (Triunfo *et al*., 2022).

In plants, the outer cell membrane has receptors which can specifically bind to chitin. Chitin-binding receptor is a protein complex, containing multiple lysin motif (LysM) domain proteins and receptor-like kinases (RLKs) or receptor-like proteins (RLPs) (Shinya *et al*., 2015). Chitin oligomers (or short-chain chitins) have high affinity to bind the receptor (Cao *et al*., 2014; Hayafune *et al*., 2014; Gubaeva *et al*., 2018). Interactions between chitin oligomers and chitin-binding receptors and downstream signaling processes may vary across different plant species but ultimately binding results in an activation of plant cellular defense response (García *et al*., 2021). The signaling process is rendered through mitogen-activated protein kinase (MAPK) cascades and in association with plant defense hormones i.e., salicylic acid, jasmonic acid and ethylene (Ramirez-Prado *et al*., 2018; Gong *et al*., 2020). On the other hand, plant receptors that can specifically bind to chitosan have not yet been reported (Yin *et al*., 2016), even though chitosan has been well described as a plant elicitor to induce plant defense (Pichyangkura and Chadchawan, 2015). The mechanism underlying chitosan-promoted plant defense is still unclear. It could be via chitin-binding receptors because partially deacetylated chitosan retains chitin-like identity and may be able to bind to the receptors. Chitosan with 60% deacetylation had higher eliciting activity than chitin oligomer (8 subunits) to enhance H_2_O_2_ production in Arabidopsis seedlings (Gubaeva *et al*., 2018). However, fully deacetylated chitosan had no effect, suggesting the acetyl group in chitin or partially deacetylated chitosan is required for chito-polymers to interact with plant receptors (Gubaeva *et al*., 2018). The chitosan used in our study was 75-85% deacetylated (Sigma-Aldrich). Hence, chitosan-induced defense responses as observed might be triggered via the binding of chitosan to chitin-binding receptors. Nevertheless, it is also feasible that the effects were induced via the binding of chitosan to its own specific receptors, which have not yet been discovered. Alternatively, chitosan may alter plant cell membrane permeability and directly signal cytoplasmic secondary messengers, such as hydrogen peroxide, nitric oxide, and phytohormones, triggering their downstream pathways (Pichyangkura and Chadchawan, 2015), or bind to chromatin in the nucleus to regulate plant defense responses (Hadwiger, 2013).

Our results also suggest that exogenous intact chitin poorly activates plant defense responses, likely due to inability to bind plant cell membrane receptors (Shinya *et al*., 2015). Based on scanning electron microscope, crude chitin aggregates into particles approximately 20 µm in diameter which is much larger than approximate size, 8 nm in diameter, of the extracellular domain of toll-like receptor, one of the classical receptors in plant and animal immune system (Fuchs *et al*., 2018). This large size mismatch and lack of available soluble chitin may impede direct binding between crude chitin and eukaryotic cell membrane receptors. Long-chain chitin nanofibers (chitin fibrils embedded in a protein matrix) can enhance cellular H_2_O_2_ production and PR gene expressions in Arabidopsis, but it is modified to be highly dispersed in water and loosely agglomerated, thereby can be easily degraded by chitinase enzymes (Egusa *et al*., 2015). Our results support the concept that natural interaction between fungal chitin and plant receptors requires digestion steps to break down polymeric cell wall chitin into soluble, recognizable chitin fragments (Shinya *et al*., 2015).

On the other hand, enhanced defense responses as observed from chitosan treatments have confirmed its eliciting properties (Chandra *et al*., 2017a; Lopez-Moya *et al*., 2021; López-Velázquez *et al*., 2021). As summarized in Fig. 9, our analyses using various techniques (including enzymatic assays, exudate proteomics and qPCR method) were well correlated, showing increased peroxidase and chitinase activities, transcript levels of defense genes in the root tissues and increased secretion of defense proteins in the exudates. Correlations between expression levels of defense genes in the tissues and their protein intensities in the exudates were also positive (Supplementary Fig. S5), implying that increased production of defense genes and proteins in tissue is correlated to secretion of the proteins into exudate. To the best of our knowledge, this is the first report showing increased secretion of defense proteins into exudate in response to chitosan. The effect appeared to be dose dependent as it was strongest in the 0.5% chitosan treatment and gradually lower in the 0.2% and 0.1% chitosan treatments. The majority of significantly increased proteins in exudate are classified as PR proteins which are usually increasingly expressed once a plant is attacked by pathogens. PR protein 1 has long been used as a genetic marker for plant systemic acquired resistance (Breen *et al*., 2017). Chitinases are key enzymes that protect plants against fungal pathogens (Vaghefi *et al*., 2013; Kaur *et al*., 2022). Thaumatin-like protein has been shown to play a role in wheat resistance against leaf rust fungus, *Puccinia triticina* (Zhang *et al*., 2018). Mulatexin protein has a strong toxicity against silkworms, helping protect mulberry tree from pest attack (Wasano *et al*., 2009). Additionally, a previous study on tomato root demonstrated that chitosan could depolarize root cell membrane and induced secretions of phytohormones, lipid signaling and phenolic compounds in root exudate (Suarez-Fernandez *et al*., 2020). Combining this information with our root exudate proteome data, it could be concluded that the impact of chitosan is not limited to plant tissues but can be extended to root metabolite and protein secretions.

**Fig. 9.**
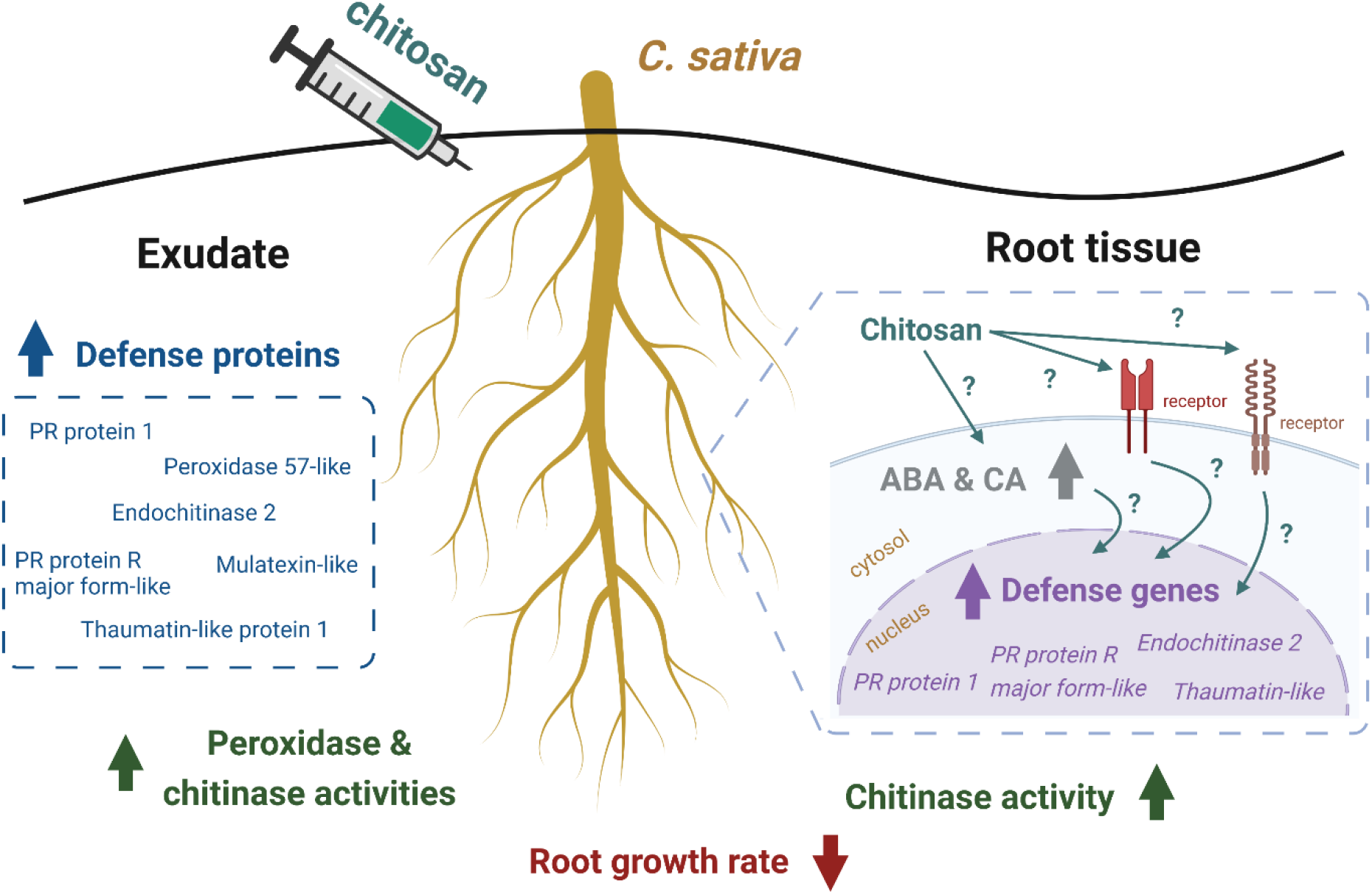
Summary of *C. sativa* root responses upon colloidal chitosan treatment as observed in this study. Root growth was affected immediately after the treatment. In the root tissue, levels of ABA hormone and CA compound were increased, defense genes were upregulated and total chitinase activity was enhanced. In the exudate, increased secretion of several defense proteins such as PR protein 1 and endochitinase 2 was detected and total peroxidase and chitinase activities were higher than control and pre-treatment. Biological mechanisms underlying root responses caused by chitosan remain uncharacterized. This figure was created with Biorender.com.

The molecular mechanism underlying chitosan promotion of plant defense is still unclear. We hypothesized that chitosan could somehow mediate biosynthesis of phytohormones, leading to a regulation of defense genes in the nucleus. In our study, phytohormone measurement showed only slight increases of ABA hormone and CA compound in the root tissue of 0.5% chitosan treatment. The levels of known defense hormones, SA and JA, were not changed in any conditions (Fig. 3). In our experiment, the sampling timepoint could be a key factor as tissue samples were collected one week after the treatment, but upregulations of genes or transcription factors related to phytohormone biosynthesis could happen within 24 hours of chitosan treatment (Colman *et al*., 2019). Higher levels of ABA detected upon chitosan treatment suggests that ABA might contribute to the increased levels of defense genes and proteins. ABA has long been proposed to associate with abiotic stress, but recently its role in relation to biotic stress has been increasingly reported (Shigenaga and Argueso, 2016). ABA signaling pathway cross talks with SA and JA defense hormone pathways, implying ABA is also involved in regulating plant disease resistance (Anderson *et al*., 2004; Berens *et al*., 2017). This is evidenced by several studies, for example, spraying 100 µM of exogenous ABA induced expression levels of PR4 and PR10 genes in lentil seedlings (Ford *et al*., 2017). Additionally, ABA hormone can modulate rice MAPK5 gene and protein biosynthesis, influencing downstream signaling processes of both disease resistance and abiotic stress tolerance (Xiong and Yang, 2003). Therefore, ABA could be one of the players, triggering defense responses in *C. sativa* root system as observed.

Moreover, we found that root growth was largely interrupted by chitosan, which is consistent with previous observations in Arabidopsis root (Lopez-Moya *et al*., 2017; Iglesias *et al*., 2019). The inhibition is due to upregulation of transcription factors involved in the auxin biosynthesis pathway, resulting in auxin accumulation at the root tips (Lopez-Moya *et al*., 2019). However, in our study auxin levels, IAA and Me-IAA, were comparable to control. In turn, higher levels of ABA and CA were detected. It is possible that auxin accumulation may take place at the early stage after chitosan exposure. The effect may ease off through time as expression level of the *YUC2* gene, encoding one of the key enzymes in the IAA biosynthesis pathway, leveled off within three days after the treatment (Lopez-Moya *et al*., 2017). Increases in cellular CA level may be another factor affecting root growth since treating plants with exogenous CA were evidenced to decrease root length, alter activity of IAA oxidase, an IAA degradation enzyme and inhibit auxin efflux (Salvador *et al*., 2013; Steenackers *et al*., 2017). Ultimately, the stalling of root growth suggests that the plant may transform energy from expanding existing roots or generating new roots to consolidate their defense system in response to chitosan. This plant adaptation process is known as growth-defense tradeoffs, which can be triggered by any abiotic and biotic factors (Huot *et al*., 2014; He *et al*., 2022). Our findings demonstrate that chitosan appears to activate this process in *C. sativa* root system. Further exploration is required to identify the underlying biological mechanisms impacting root growth and causing upregulation of defense responses. The knowledge gained would benefit crop improvement to maximize crop yield with a balance of disease resistance (Silva *et al*., 2019). After all, chitosan could be a potential elicitor for agricultural application, especially in hydroponic setup to counteract fungal pathogen attacks under safe and practical conditions.

## Conclusion

Finding safe and effective solutions to manage crop diseases is an essential task to support *C. sativa* production. Chitin and chitosan are natural elicitors, known to promote plant defense. They may be similar in terms of structure, but their effects on plant cell recognition, physiological responses and molecular processes are highly distinct. We found that colloidal chitin has very low impact on *C. sativa* defense promotion while colloidal chitosan can enhance defense responses in root tissue and exudate. The key finding was the detection of several defense proteins including PR proteins, chitinases and thaumatin-like proteins that were increasingly secreted upon chitosan treatment. This was confirmed by increases in total activities of peroxidase and chitinase enzymes in the exudate. However, root growth was interrupted after chitosan exposure. Biological pathways underlying defense promotion, but root growth inhibition caused by chitosan remain uncharacterized. Increased cellular levels of ABA and CA were detected and could be one of the underlying factors. Further study is required to investigate how the plant recognizes the chitosan molecules and what signaling pathways lead to the root transformation from growth to defense. Nonetheless, chitosan has potential for implementation in *C. sativa* production, particularly in hydroponic cultivation to manage waterborne fungal diseases.

## Abbreviations

ABA: abscisic acid
ACN: acetonitrile
CA: cinnamic acid
ESI: electrospray ionization
FW: fresh weight
*N*-GlcNAc: *N*-acetyl-D-glucosamine
IAA: indole-3-acetic acid
JA: jasmonic acid
LC: liquid chromatography
LFQ: label-free quantification
MAPK: mitogen-activated protein kinase
MS: mass spectrometry
MWCO: molecular weight cutoff
OPDA: 12-oxo-phytodienoic acid
PCA: principle component analysis
PR: pathogenesis-related
qPCR: quantitative real-time PCR
Root-TRAPR: Root-Transparent Reusable Affordable three-dimensional Printed Rhizo-hydroponic
RSA: root surface area
SA: salicylic acid
SQRT: square root
TEAB: triethylammonium bicarbonate
TFA: trifluoroacetic acid
THC: tetrahydrocannabinol

## Supplementary data

The following supplementary data are available at JXB online

**Fig. S1** Root growth rates measured from root length and surface area.

**Fig. S2** Phytohormone levels of IAA, methyl-IAA and zeatin.

**Fig. S3** Additional PCA plots of exudate proteomes.

**Fig. S4** Dose response curves comparing qPCR transcript levels and chitosan concentrations of four defense-related genes.

**Fig. S5** Correlation plots between qPCR transcript levels and exudate protein intensities of four studied proteins.

**Fig. S6** Correlation plots of endochitinase 2 transcript level and total chitinase activities in root tissue and exudate.

**Table S1** Protein identification details and statistical analysis of exudate proteomes.

**Table S2** Details of genes and primers used in the qPCR analysis.

**Table S3** Extended function prediction scores of 57 identified exudate proteins.

## Acknowledgements

We thank David Brian (Southern Hemp Co.) for supplying hemp seeds. We thank 3D printing teams at the New Experimental Technology Lab (NExT Lab), Melbourne School of Design and the Telstra Creator Space, Faculty of Engineering and Information Technology, University of Melbourne for printing the Root-TRAPR frames. We thank the Engineering Workshop, University of Melbourne for making the Root-TRAPR acrylic sheets. We thank Swati Varshney and Nicholas Williamson at the Mass Spectrometry and Proteomics Facility (MSPF), Bio21 Molecular Science and Biotechnology Institute, University of Melbourne for advice and support on proteomic analysis. We thank Tannaz Zare and Jacob Calabria for technical guidance on RNA extraction and qPCR analysis.

## Author contributions

PS, AI, JSP, RW and BAB designed the study. PS conducted the experiments and analyzed the data. SN performed proteomics analysis. AI, JSP, RW and BAB provided guidance and technical support throughout the study. PS prepared the manuscript. All authors edited and approved the final version.

## Data availability

The data that support the findings of this study are available in the supplementary materials published online. Raw data of root parameters and all biochemical analyses will be uploaded to Dyrad online repository and raw proteomics data will be deposited to Proteomics Identification Database (PRIDE) upon revision process.

