## Supplementary Fig. S1-S6 for "Effects of chitin and chitosan on root growth, biochemical defense response and exudate proteome of *Cannabis sativa*"

### Supplementary data

The following supplementary data are available at JXB online

**Fig. S1** Root growth rates measured from root length and surface area.

**Fig. S2** Phytohormone levels of IAA, methyl-IAA and zeatin.

**Fig. S3** Additional PCA plots of exudate proteomes.

**Fig. S4** Dose response curves comparing qPCR transcript levels and chitosan concentrations of four defense-related genes.

**Fig. S5** Correlation plots between qPCR transcript levels and exudate protein intensities of four studied proteins.

**Fig. S6** Correlation plots of endochitinase 2 transcript level and total chitinase activities in root tissue and exudate.

**Table S1** Protein identification details and statistical analysis of exudate proteomes.

**Table S2** Details of genes and primers used in the qPCR analysis.

**Table S3** Extended function prediction scores of 57 identified exudate proteins.

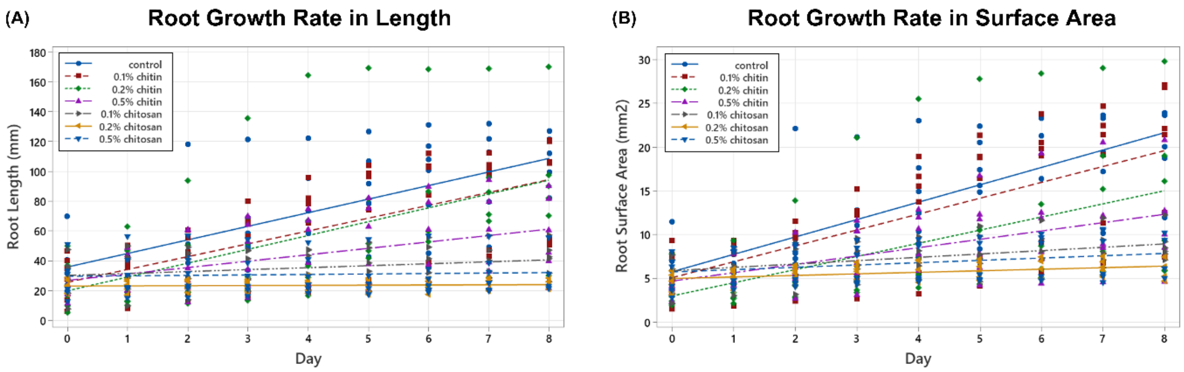


| Condition | Linear regression | *p*-value |
|---|---|---|
| Control | Y = 35.95 + 9.10X | - |
| 0.1% chitin | Y = 25.58 + 8.61X | 0.778 |
| 0.2% chitin | Y = 19.87 + 9.28X | 0.920 |
| 0.5% chitin | Y = 26.87 + 4.30X | 0.007 |
| 0.1% chitosan | Y = 30.18 + 1.31X | 0.000 |
| 0.2% chitosan | Y = 23.22 + 0.11X | 0.000 |
| 0.5% chitosan | Y = 29.62 + 0.31X | 0.000 |

| Condition | Linear regression | *p*-value |
|---|---|---|
| Control | Y = 5.77 + 1.983X | - |
| 0.1% chitin | Y = 5.08 + 1.814X | 0.586 |
| 0.2% chitin | Y = 2.99 + 1.501X | 0.124 |
| 0.5% chitin | Y = 4.69 + 0.954X | 0.001 |
| 0.1% chitosan | Y = 5.87 + 0.382X | 0.000 |
| 0.2% chitosan | Y = 5.00 + 0.177X | 0.000 |
| 0.5% chitosan | Y = 5.73 + 0.264X | 0.000 |

**Fig. S1** Linear regressions of root growth rate measured from root length (A) and root surface area (B) from treatment day (day 0) until sample collection (day 8) among control, chitin and chitosan treatments within six biological replicates. Regression coefficient *p*-value was analyzed from an interaction between treatment and day. The comparison was performed between respective treatment with control. Significant differences ($P < 0.05$) were observed from 0.5% chitin and 0.1-0.5% chitosan treatments.

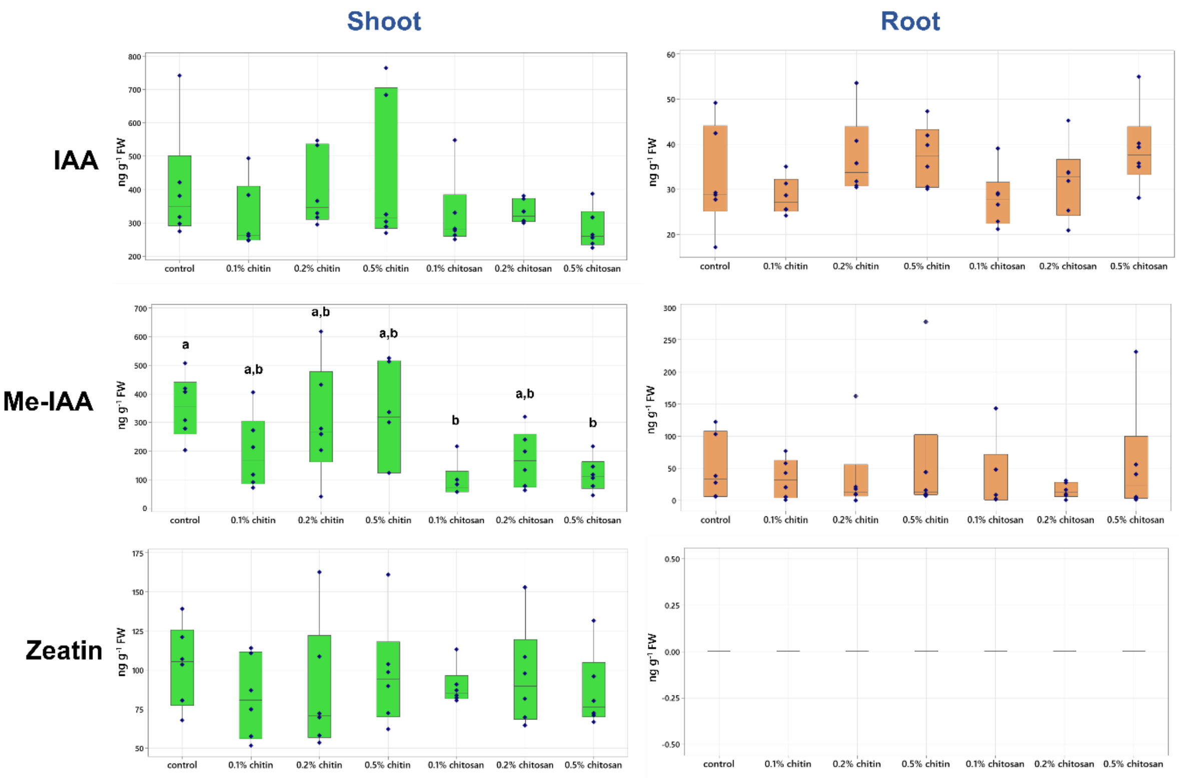


**Fig. S2** Phytohormone levels of indole-3-acetic acid (IAA), methyl-IAA (Me-IAA) and zeatin measured from shoot and root tissues of control, chitin and chitosan treatments within six biological replicates. Letters (a-b) refer to statistically significant difference ($P$ < 0.05) using one-way ANOVA, followed by Tukey's *post hoc* analysis.

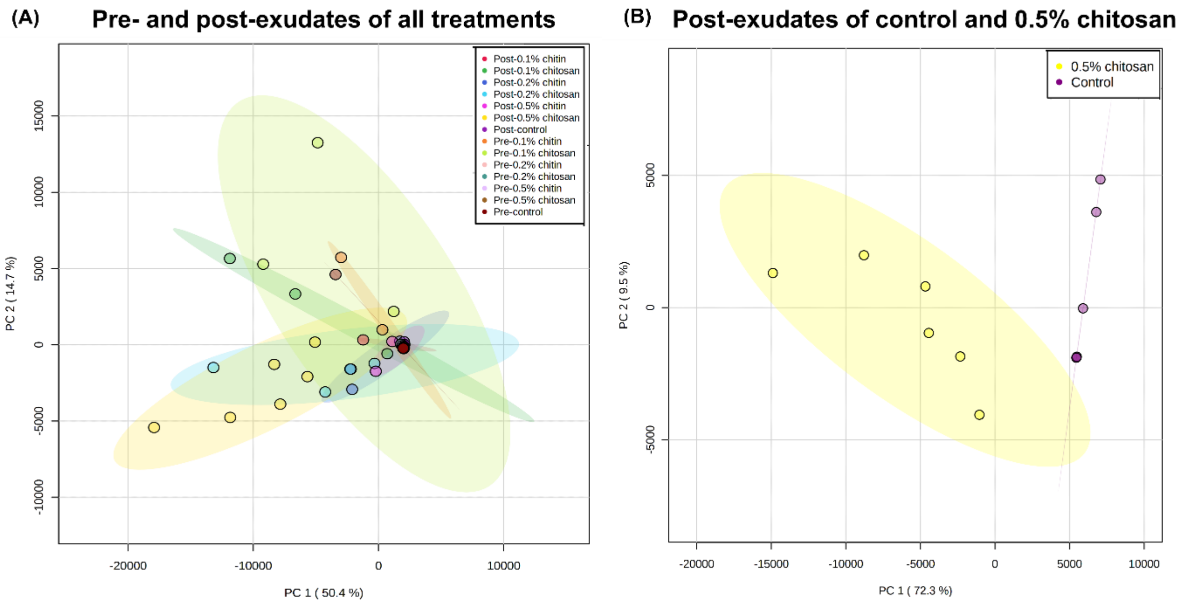


**Fig. S3** PCA plots of exudate proteomes across pre- and post-exudate of all samples (A) and only post-exudate proteomes of control and 0.5% chitosan treatment (B). Overview PCA shows majority of the samples were clustered near the zero origin with few samples scattered away (A). After treatment, a clear separation between post-exudates of control and 0.5% chitosan treatment was detected (B). Six biological replicates were analyzed per treatment.

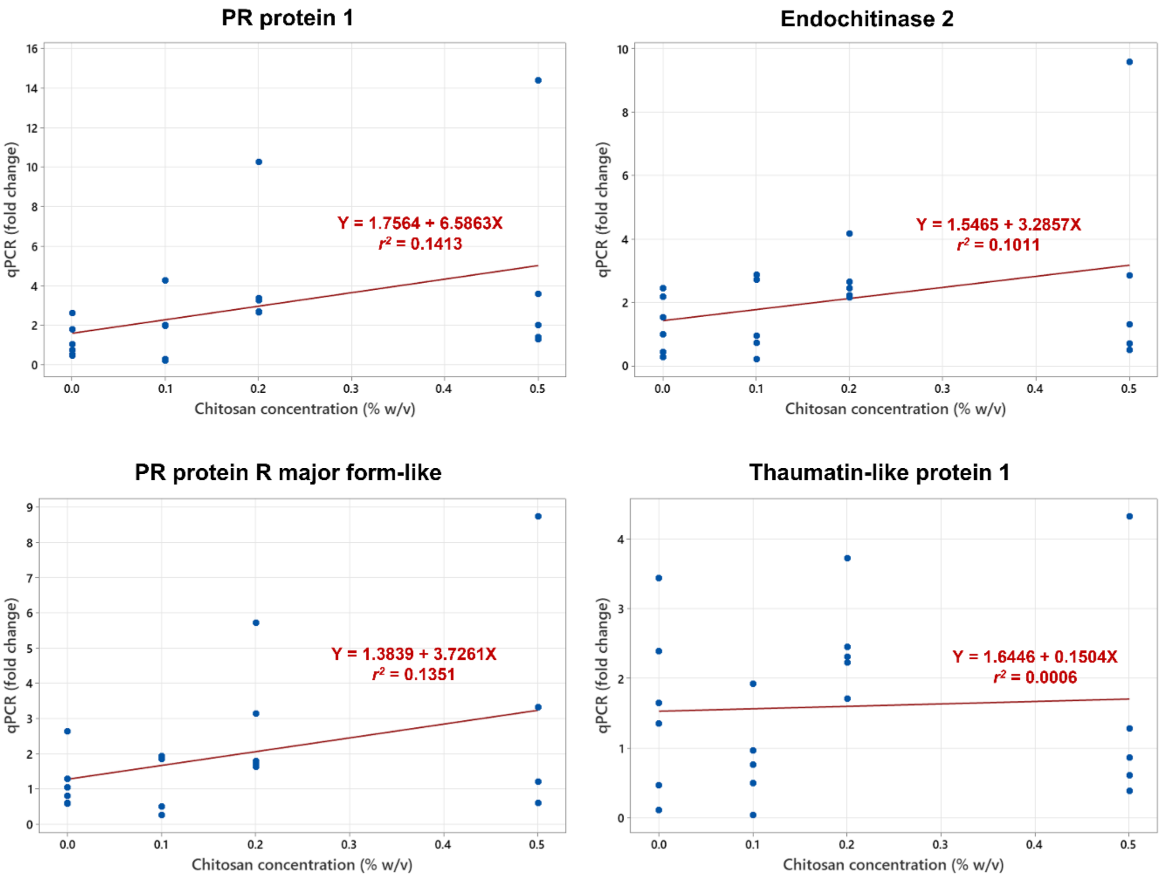


**Fig. S4** Dose response curves comparing between qPCR transcript levels of PR protein 1, endochitinase 2, PR protein R major form-like and thaumatin-like protein 1 and chitosan concentrations of 0% (control), 0.1%, 0.2% and 0.5% w/v within five biological replicates. Positive relationships were detected from PR protein 1, endochitinase 2 and PR protein R major form-like.

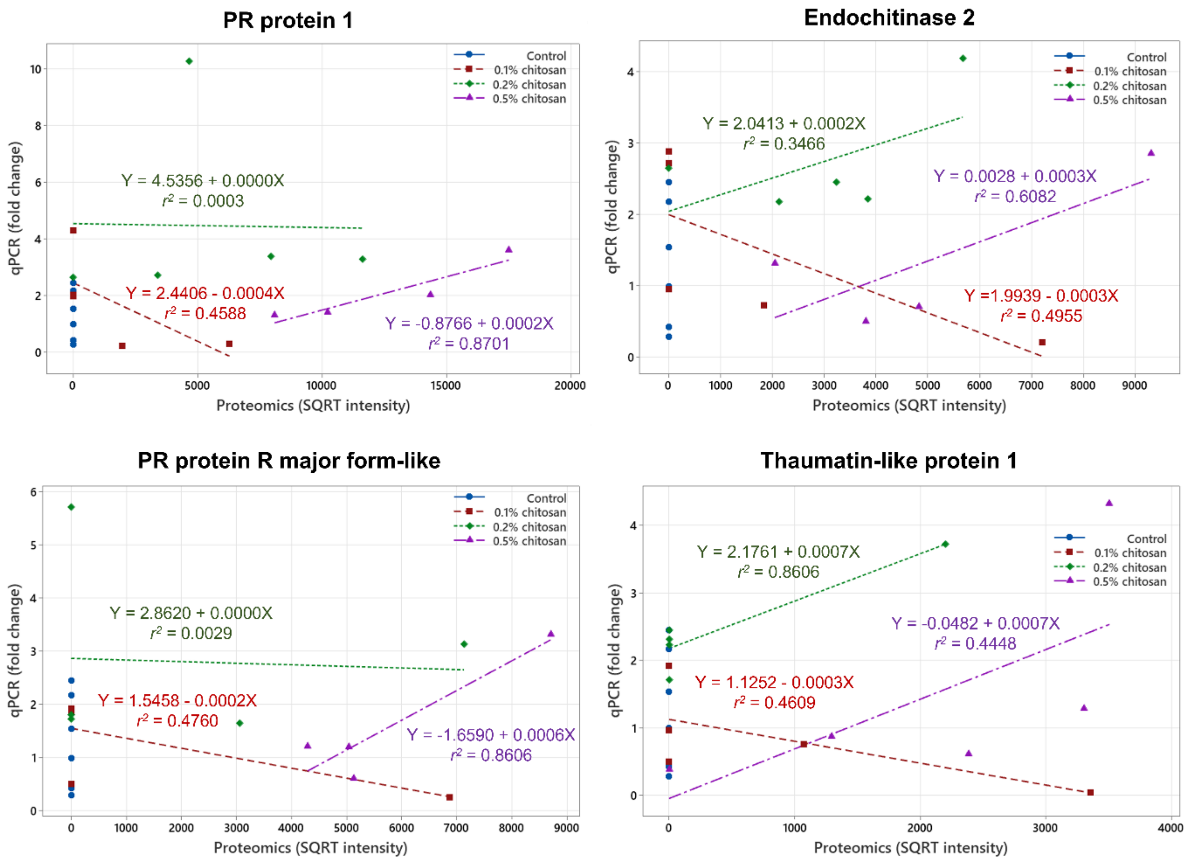


**Fig. S5** Correlation plots between qPCR transcript levels and exudate proteomics intensities of PR protein 1, endochitinase 2, PR protein R major form-like and thaumatin-like protein 1 within five biological replicates. Regression equation and coefficient of determination ($r^2$) value are indicated next to the regression line. Positive correlations were observed from 0.5% chitosan treatment of all four proteins and 0.2% chitosan treatment of endochitinase 2 and thaumatin-like protein 1.

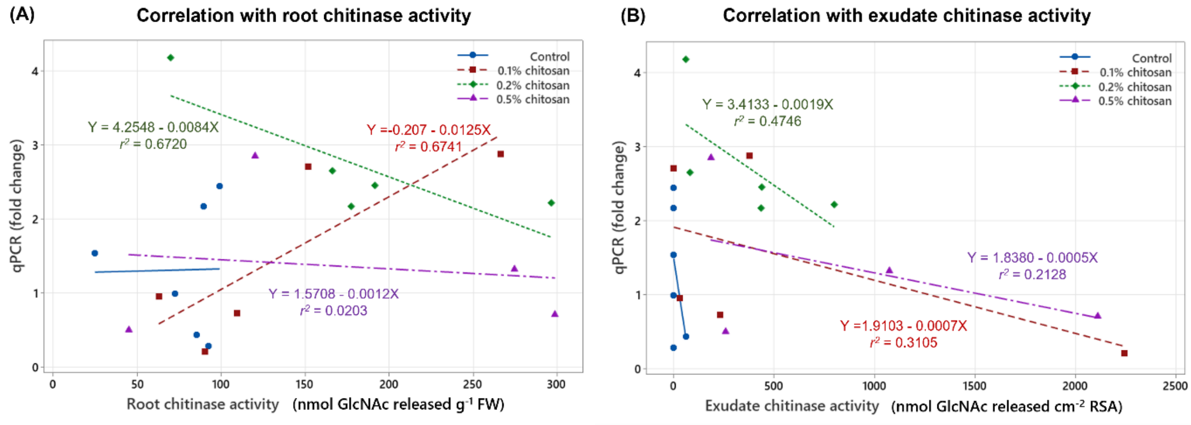


**Fig. S6** Correlation plots between endochitinase 2 transcript level and total chitinase activities measured from root tissues (A) and exudates (B) of control and chitosan treatments within five biological replicates. Regression equation and coefficient of determination ($r^2$) value are indicated next to the regression line.

**Table S1** Protein identification details and statistical analysis of all 57 proteins detected in the exudates.

>> Supplied in Excel file

**Table S2** Details of genes and primers used in the qPCR analysis.

>> Supplied in Excel file

**Table S3** Extended prediction scores of cellular location, molecular function and biological process of all 57 proteins detected from the exudates.

>> Supplied in Excel file
